# Shaping cryogenic 3D volume imaging by pFIB ion species and milling voltage

**DOI:** 10.64898/2026.09.16.751696

**Authors:** Sanja Sviben, Christopher Thompson, Corinna Braun, Krishnan Venkataraman, Jessica Heebner, Ron Kelley, M. Zuhaib Qayyum, Abhay Kotecha, Ali Khalighifar, Georgios Skiniotis

## Abstract

Cryogenic plasma focused ion beam-scanning electron microscopy (cryo-pFIB-SEM) enables nanoscale volume imaging of vitrified cells and tissues, but the fidelity of each newly exposed block-face depends on how ions interact with compositionally heterogeneous biological material. Using xenon, argon, oxygen, and nitrogen plasma beams across a 30-2 kV voltage range, we examined how ion species and accelerating voltage influence block-face quality and ultrastructural fidelity. Curtaining imposed a species-dependent lower limit on accelerating voltage, with xenon and argon supporting uniform milling to 10 kV and oxygen and nitrogen to 15 kV. Monte Carlo simulations predicted substantially reduced ion penetration and atomic displacement under these species-specific low-kV conditions. Experimentally, lowering milling voltage suppressed charging and topographic artifacts, with the largest effects observed at heterogeneous interfaces. Oxygen and nitrogen produced greater membrane sharpness and organelle contrast than the noble gases, and these differences influenced machine-learning-based recognition of fine organelle features in 3D datasets. Together, these findings show that cryo-pFIB-SEM image quality cannot be optimized by accelerating voltage or ion species independently. Instead, milling performance reflects a tradeoff among surface fidelity, charging, contrast, and curtaining that depends on local specimen composition. Species-specific low-kV milling therefore provides a practical framework for matching cryo-pFIB-SEM acquisition conditions to the structural and biophysical properties of the biological target.

## Introduction

Cryogenic focused ion beam-scanning electron microscopy (cryo-FIB-SEM) has become an important tool in structural cell biology, both for preparing electron-transparent lamellae for cryo-electron tomography (cryo-ET) and for three-dimensional (3D) volume imaging of vitrified cells and tissues^1–3^. In volume imaging, iterative ion-beam milling and SEM imaging are used to reconstruct cellular volumes at nanometer scale resolution, providing organelle-level context that complements the molecular information accessible by cryo-ET^4,5^. Unlike conventional FIB-SEM of fixed, stained, resin-embedded specimens^6,7^, cryogenic volume imaging interrogates unstained heterogeneous biological material in which image contrast arises from intrinsic differences in composition, surface topography, and charge distribution^4,8,9^. The freshly exposed block-face contains interfaces among aqueous cytoplasm, protein-dense structures, membranes and lipid-rich compartments, each of which may respond differently to ion bombardment. Milling thus not only exposes the surface to be imaged but can also alter its physical properties, thereby influencing the biological information recorded in each SEM image.

High-energy ions interact with vitrified biological material through elastic and inelastic scattering, generating collision cascades, atomic displacements, sputtering, local heating, and ion implantation. These interactions can modify the newly exposed block-face, influencing charging, curtaining, contrast, and surface morphology^10,11^. Studies have begun to investigate ion induced damage in cryo-FIB generated lamellae for cryo-ET, where ion penetration under the surface can damage the specimen and can be reduced by lowering the ion-beam accelerating voltage^12–14^. Cryo-FIB-SEM volume imaging presents a fundamentally different problem, because each milling step creates a new surface, the block-face, that is immediately imaged by SEM. Surface morphology, charging, secondary-electron (SE) contrast, and milling across different compositional interfaces are therefore critical determinants of image quality^4,8^. Lowering the ion-beam voltage might improve block-face fidelity by reducing ion-induced surface modification. On the other hand, lower ion-beam voltage can reduce sputtering efficiency and increase curtaining. These competing effects are expected to depend on ion species, whose mass and reactivity influence ion penetration, sputtering, and interactions with heterogeneous biological material^9,15^. Thus, the effects of milling voltage and ion species must be considered together rather than as independent parameters, an interplay that has not been systematically explored in cryo-FIB-SEM volume imaging.

Here, we examine how plasma ion species and accelerating voltage jointly determine block-face quality in cryogenic volume imaging using xenon, argon, oxygen, and nitrogen beams across a 30-2 kV voltage range. We first establish species-specific lower voltage limits and compare these conditions with the conventional 30 kV milling. We then combine Monte Carlo simulations with quantitative analysis of SEM resolution, contrast, sharpness, charging, and block-face topography to determine how these parameters affect the resulting images. Finally, we assess whether these differences extend to 3D recognition of organelle ultrastructure using machine-learning (ML)-based segmentation. We show that lowering milling voltage reduces predicted ion-induced damage and suppresses prominent charging and topographic artifacts at compositionally heterogeneous interfaces without compromising the spatial information recovered by SEM imaging. At the same time, ion species impose distinct tradeoffs: oxygen and nitrogen produce sharper membrane boundaries and greater organelle contrast but are more susceptible to curtaining, whereas xenon and argon tolerate lower accelerating voltages. Together, these findings establish that cryo-pFIB-SEM image quality emerges from the interaction among ion species, accelerating voltage, and local specimen composition, providing a basis for selecting milling conditions according to the structural and biophysical properties of the biological target.

## Results

### Ion species defines the minimum milling voltage for cryogenic block-face imaging

We first sought to establish the lowest accelerating voltage compatible with uniform block-face imaging for each plasma ion species. Under constant SEM imaging conditions and low ion-beam currents^9^ we acquired SEM micrographs of vitrified mouse epithelial cells following xenon, argon, oxygen, or nitrogen milling across a voltage series of 30, 20, 15, 10, 8, 5, and 2 kV (Fig. 1a, Supplementary Table 1). As the accelerating voltage was reduced, curtaining artifacts emerged, initially in the lower portion of the milled surface, and block-face uniformity progressively deteriorated. To quantify this effect, we implemented a curtaining scoring algorithm based on the spatial frequency and amplitude of intensity variations along the milling direction (Figs. 1e,f).

**Figure 1.**
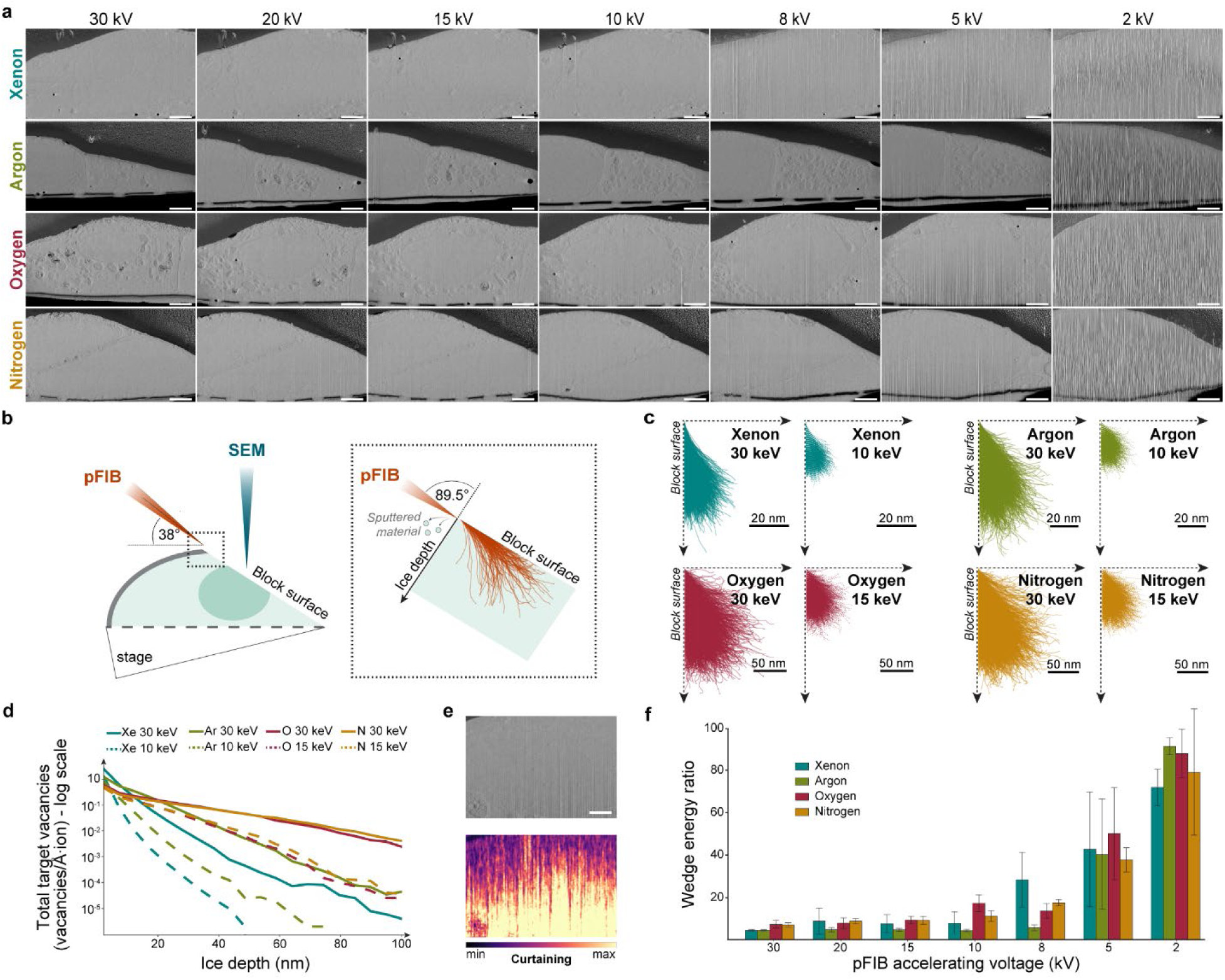
Determining low voltage pFIB milling regimes for cryogenic volume imaging. (A) A series of block-face SEM micrographs of vitrified cells milled with different ion species at different accelerating voltages. Scale bar: 1 µm. (B) Schematic of the milling and imaging geometry used in this study: a pre-tilted shuttle positions the FIB at a 38° incidence angle to the grid plane, with SEM imaging performed at a 4 mm working distance. Inset (black dotted rectangle) depicts the geometry used for the Monte Carlo simulations in (C) corresponding to an ion beam incidence angle of 89.5° relative to the milled sample surface. (C) Monte Carlo simulations of damage depth profiles for each ion species at 30 keV and the chosen low-keV milling regime, showing trajectories and implantation depth of 10 000 ions in amorphous ice (density 0.94 g/cm^3^). (D) Depth profile of total target vacancies from Monte Carlo simulations for different ion species at 30 keV and the corresponding low-keV condition. Depth (x-axis) represents distance into the amorphous ice target from the block-face. (E) SEM micrograph after argon milling at 5 kV, showing curtaining artifacts propagating from the bottom to the top of the block-face, with the corresponding curtaining heatmap. Scale bar: 1 µm. (F) Quantification of curtaining artifacts across three biological replicates, shown for each ion species across high and low accelerating voltages.

The voltage dependence of curtaining differed substantially among ion species. Xenon and argon maintained relatively low curtaining scores down to 10 kV, below which curtaining increased. Oxygen and nitrogen exhibited higher curtaining across the voltage series and became unsuitable for uniform block-face preparation below 15 kV. We therefore defined 10 kV as the lowest practical condition for xenon and argon and 15 kV for oxygen and nitrogen. These species-specific low-kV conditions, together with 30 kV as a reference condition, were used for subsequent comparisons.

### Simulations predict low-kV milling reduces ion penetration and displacement damage

To establish a physical framework for comparing the standard 30 kV condition with the species-specific low-kV regimes defined above, we performed Monte Carlo simulations of ion trajectories in amorphous ice as a proxy for vitrified biological material (Figs. 1b,c). For all four ion species, reducing beam energy resulted in substantially shallower ion trajectories. At 30 keV, xenon exhibited the shortest projected range, followed by argon, whereas nitrogen penetrated most deeply. This relative ordering was maintained under the corresponding low-keV conditions, although the predicted penetration ranges were compressed for all four species.

We next examined target vacancy profiles as a spatial measure of atomic displacement within the simulated specimen (Fig. 1d). Reducing ion energy decreased both the depth over which vacancies were generated and the total number of vacancies. Near the block surface, xenon generated the highest surface vacancy density per incident ion under both energy regimes, whereas nitrogen generated fewer vacancies despite its greater penetration depth. Thus, the simulations predict that reducing accelerating voltage from 30 kV to the species-specific low-kV conditions decreases both the spatial extent and magnitude of ion-induced atomic displacement.

### Reducing milling voltage preserves block-face SEM resolution

Having established that lower accelerating voltage is predicted to reduce ion penetration and displacement damage, we next asked whether this reduction in ion energy compromises the spatial information recovered by SEM. To minimize variability arising from differences among cells and cellular regions, we used a feature-to-feature comparison workflow in which micrographs were acquired from the same region sequentially before and after switching from 30 kV to the corresponding species-specific low-kV condition (Figs. 2a,b and Supplementary Fig. 1). We performed Fourier ring correlation (FRC) analysis on spatially distributed image patches across the cell and calculated the mean FRC-derived resolution for each dataset (Fig. 2c and Supplementary Figs. 2 and 3a).

**Figure 2.**
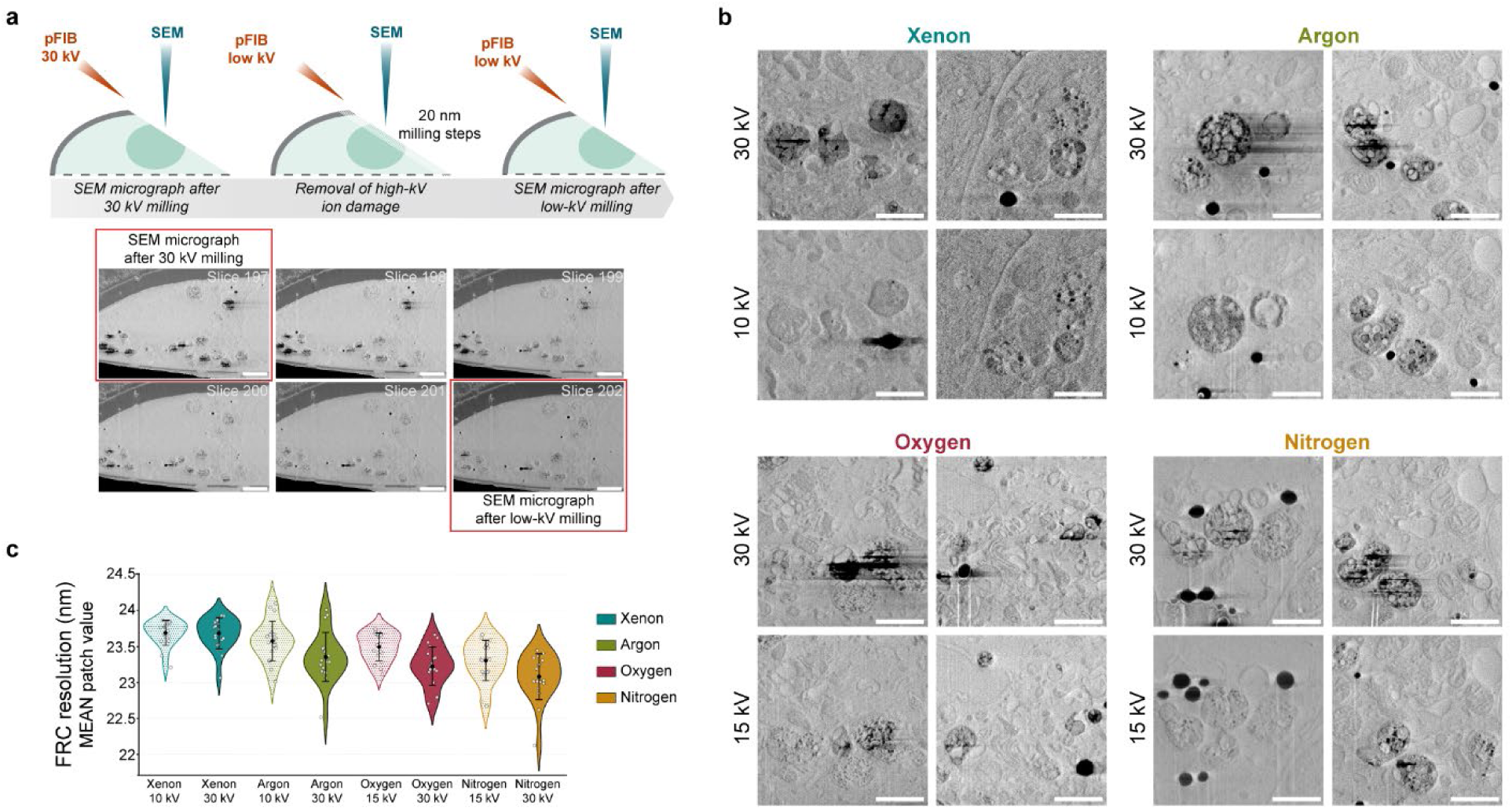
The effect of pFIB milling on SEM block-face imaging. (A) Schematic of the ‘feature-to-feature’ workflow used to assess the effects of 30 kV and low-kV pFIB milling on the same ultrastructural features. Following block-face preparation with pFIB at 30 kV and initial SEM imaging, the pFIB was switched to the selected low-kV regime, and 100 nm of material was removed in 20 nm increments before acquiring a second SEM micrograph for comparison. Representative SEM micrographs over the course of the workflow are shown below. Scale bar: 2 µm. (B) Representative SEM micrographs showing changes in SEM image quality following high-versus low-voltage pFIB milling. SEM imaging conditions were held constant across all experiments. Scale bar: 500 nm. (C) Fourier Ring Correlation (FRC) analysis based on mean patch value for whole-cell SEM micrographs from the ‘feature-to-feature’ workflow across five biological replicates, each with up to five technical replicates. Solid violin plots represent 30 kV milling regimes; dotted violin plots represent low-kV milling regimes, color-coded by ion species.

Across all four ion species, reducing the milling voltage to the species-specific low-kV condition did not produce a meaningful change in mean patch resolution compared with 30 kV. Minimum and maximum patch-resolution values, representing the highest- and lowest-resolution regions, respectively, were similarly distributed across ion species and milling voltages (Supplementary Fig. 3). These results indicate that reducing the milling voltage within the usable range does not substantially compromise the spatial resolution of block-face SEM imaging.

### Ion species and milling voltage differentially affect organelle image quality

Because global spatial resolution was largely preserved, we next examined image properties more directly related to the resolvability of cellular structures. Using the feature-to-feature comparison workflow, we quantified charging, sharpness, and contrast both at the whole-cell level and within individual organelle classes (Fig. 3).

**Figure 3.**
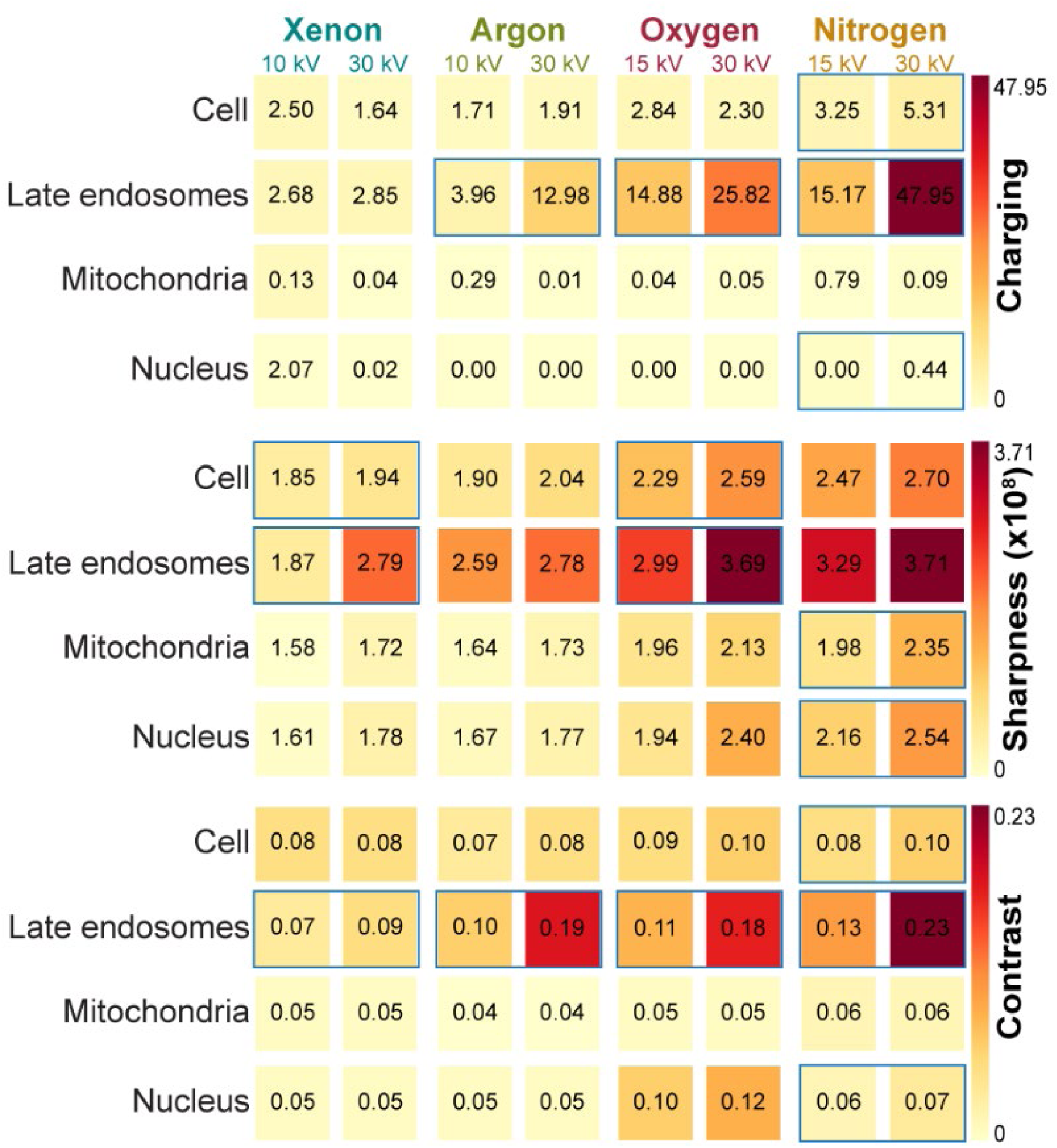
SEM micrograph quality quantification in relation to pFIB ion species and milling voltage. Heatmaps show charging fluctuations, sharpness, and contrast metrics at both the whole-cell and organelle level across five biological replicates, each with up to five technical replicates. Values were normalized across all ion species and milling voltages tested. Blue frame rectangles mark 30 kV-low kV pairs with statistical significance (p = 0.0625).

At the whole-cell level, differences between 30 kV and species-specific low-kV conditions were generally modest. Nitrogen showed consistent replicate-level reductions in charging and contrast at lower voltage, while oxygen and xenon showed consistent replicate-level reductions in sharpness. These effects reached the strongest statistical support possible for a two-sided Wilcoxon signed-rank test with five paired biological replicates (p = 0.0625). Nevertheless, ion-species-dependent differences were apparent across both voltage regimes. In particular, oxygen- and nitrogen-milled surfaces consistently exhibited higher sharpness than xenon- and argon-milled surfaces, indicating that ion species influences the definition of ultrastructural boundaries independently of accelerating voltage.

Analysis at the organelle level revealed effects that were less apparent in whole-cell measurements. Late endosomes were particularly sensitive to both ion species and accelerating voltage. Contrast within late endosomes was highest following nitrogen milling, while charging was reduced under the species-specific low-kV conditions, with the most pronounced reductions observed for oxygen and nitrogen. In regions affected by charging at 30 kV, low-kV milling restored membrane visibility and made these structures readily traceable for subsequent segmentation (Fig. 2b and Supplementary Fig. 1). Thus, the effects of milling conditions were more apparent at the level of individual organelles than in whole-cell measurements, particularly within compositionally heterogeneous and lipid-rich compartments.

### Low-kV milling suppresses topographic artifacts at compositionally heterogeneous interfaces

We next examined whether the observed changes in image quality were associated with physical alterations of the block-face. Following 30 kV milling, SEM images revealed a prominent bright-line feature at the interface between the GIS platinum protection layer and the cell surface. This feature was largely absent from backscatter-electron images, with only modestly increased backscatter signal along parts of the interface for argon and nitrogen (Supplementary Fig. 4). Similar features were observed at interfaces between materials likely to differ in sputtering behavior, particularly between lipid-rich regions and cytosol, including beneath the carbon support film and at the boundaries of lipid droplets and lipid rich domains within late endosomes (Fig. 4a). This was observed with all four ion species and diminished progressively as the milling voltage was reduced (Supplementary Fig. 5).

**Figure 4.**
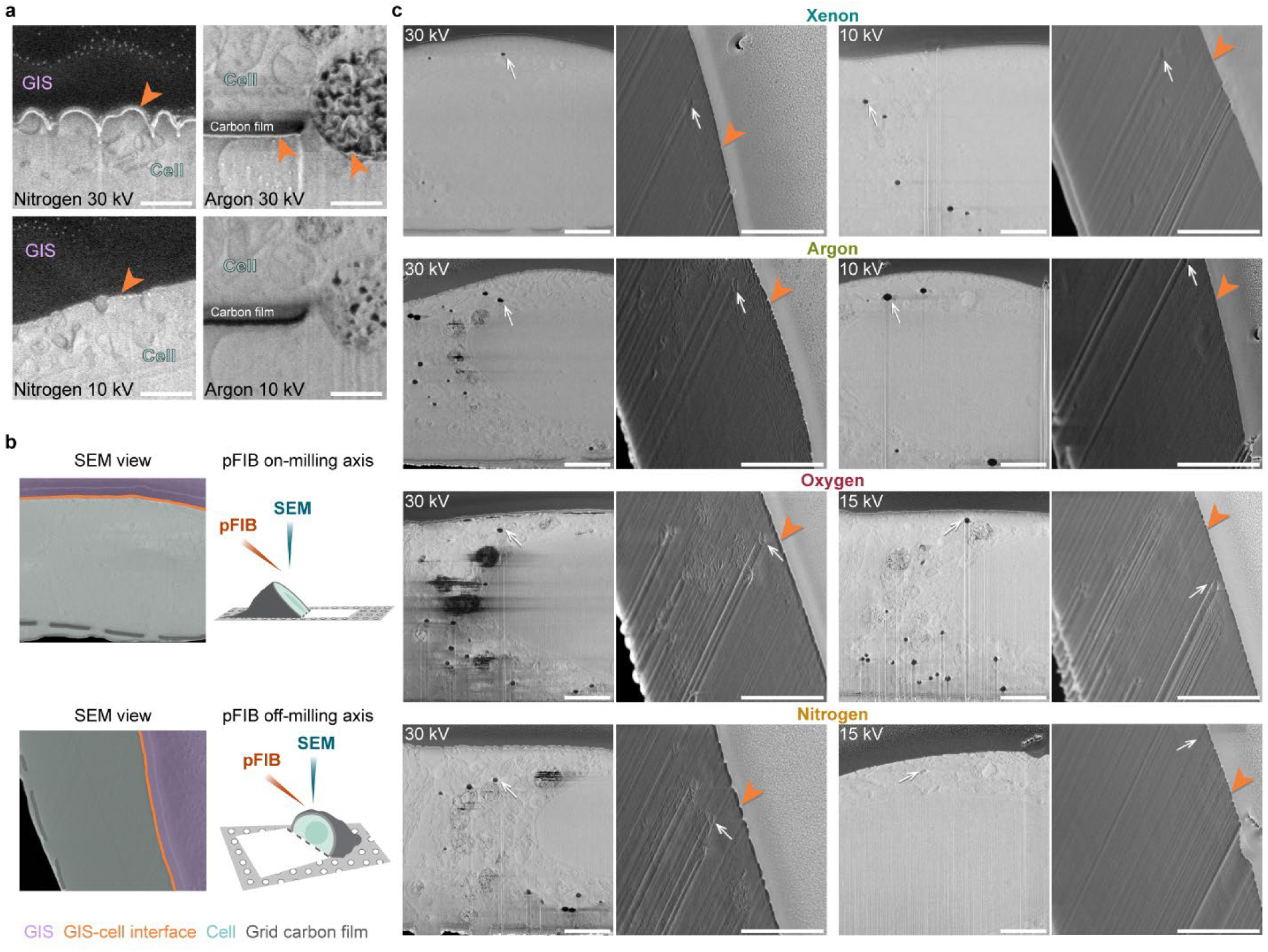
Topographic artifacts at compositional interfaces on the SEM block-face. (A) Representative SEM micrographs showing artifacts following 30 kV milling. A bright-line feature (orange arrowhead) is visible at several compositional interfaces: the GIS (gas injection system) platinum protection layer-cell surface, the EM grid carbon film-cell interface, and below lipid-rich densities. The artifact is absent or substantially suppressed after low-kV milling. Scale bar: 500 nm. (B) Sample orientation in respect to pFIB and SEM beams used to assess topographic artifacts in (C). On-axis and off-axis imaging refers to block-face position in relation to initial pFIB milling axis. (C) Representative SEM micrographs showing block-face topography under on-axis and off-axis imaging, as a function of milling ion species and voltage. Paired images are marked with landmarks denoting the same cellular feature (white arrow) and highlighting an undercut at the GIS - cell interface (orange arrowhead) that is reduced under low-kV conditions. Scale bar: 1 µm.

To determine whether the bright-line feature corresponded to a physical alteration of the block-face, we acquired SEM images after rotating the specimen away from the initial milling orientation (Supplementary Table 2). Off-axis imaging revealed an indentation or undercut at the GIS-cell interface that spatially coincided with the bright-line artifact observed in on-axis images (Figs. 4b,c). This feature was reproducibly observed following 30 kV milling with all four ion species and was substantially reduced at lower milling voltage, linking the image artifact to voltage-dependent modification of block-face topography. On the other hand, reducing the pFIB current at 30 kV did not improve the block-face appearance, indicating that the effect was not primarily current-dependent (Supplementary Fig. 6).

Off-axis imaging also revealed ion-species- and voltage-dependent differences in surface texture. Membranes and other lipid-rich boundaries appeared selectively accentuated following 30 kV milling with argon, oxygen, and nitrogen, producing an etched appearance that was reduced under the corresponding low-kV conditions. These observations demonstrate that both accelerating voltage and ion species influence the surface topography generated at compositionally heterogeneous interfaces and provide a structural correlate for the species-dependent differences in membrane sharpness and contrast observed in block-face SEM images.

### Milling conditions determine ML-based recognition of organelle ultrastructure

We next asked whether the differences observed in 2D image quality translated into the recognition of ultrastructural features in 3D volume datasets. We therefore performed ML-based segmentation of high- and low-kV slice-and-view datasets (Supplementary Videos 1–16). Segmentation outputs were minimally processed to remove small, isolated pixel islands, without addition of manual labels (Fig. 5 and Supplementary Fig. 7).

**Figure 5.**
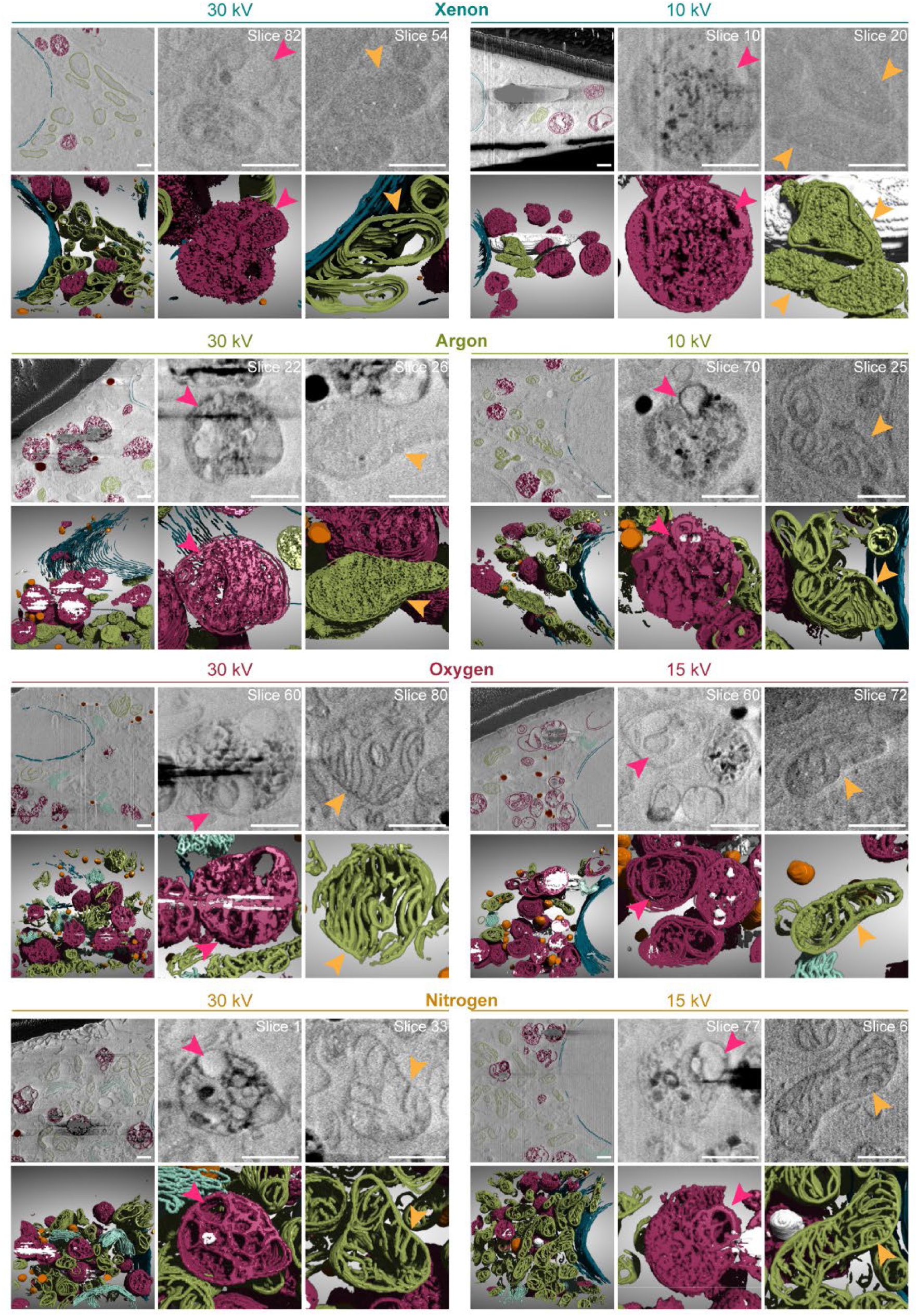
Machine learning-based segmentation and 3D visualization of cellular compartments in volume imaging experiments using different plasma gas species and milling voltages. Dark magenta indicates membranes in late endosomes, green indicates membranes associated with mitochondria, dark blue indicates the nuclear membrane, light blue indicates Golgi apparatus, orange indicates lipid bodies and white indicates charged regions. Magenta arrowheads label corresponding regions of late endosomes in SEM micrographs and their accompanying 3D visualizations, while orange arrowheads label corresponding regions of mitochondria in SEM micrographs and accompanying 3D visualizations. Scale bar: 500nm.

The extent of ultrastructural recognition varied substantially among ion species and milling conditions. Xenon produced the least defined intraluminal ultrastructure. At 30 kV, segmentation primarily identified organelle boundaries, while late-endosomal internal densities remained poorly resolved. At 10 kV, additional mitochondrial and endosomal features became detectable, although many organelles remained unrecognized.

Argon at 30 kV produced a similar degree of structural definition to xenon at 10 kV, whereas argon at 10 kV enabled segmentation of mitochondrial cristae and distinct late-endosomal intraluminal densities. Oxygen and nitrogen provided greater structural definition at 30 kV than xenon or argon at the same voltage, enabling segmentation of mitochondrial cristae and late-endosomal intraluminal densities. Although charging was more pronounced under these conditions, it decreased at lower voltage while fine ultrastructural features remained well defined. Golgi cisternae were sufficiently resolved for segmentation in nitrogen datasets acquired at both 30 and 15 kV, in oxygen datasets acquired at 15 kV, and in argon datasets acquired at 10 kV (Supplementary Fig. 8).

These results show that the voltage required to reveal fine organelle ultrastructure depends strongly on ion species. Lower-kV milling generally improved the visibility of internal features, particularly in datasets affected by charging, while the reactive oxygen and nitrogen plasma ion species provided greater structural definition overall.

### SEM spatial resolution remains stable during automated low-kV volume acquisition

We next determined whether SEM spatial resolution remained stable during automated volume acquisition. Patch-based FRC-derived resolution was monitored across consecutive block-face images acquired at 30 kV and under the corresponding species-specific low-kV conditions (Supplementary Figs. 9 and 10). Although mean resolution was comparable between voltage regimes, its temporal behavior differed significantly (image number × voltage interaction, p = 0.006). Across ion species, resolution deteriorated by approximately 1.04 nm per 100 images at 30 kV (95% CI, 0.49-1.60), whereas no systematic change was detected under the species-specific low-kV conditions (−0.07 nm per 100 images; 95% CI, −0.64-0.50). Temporal trends did not differ detectably among ion species (p = 0.109). Thus, low-kV milling preserved mean SEM spatial resolution and was associated with greater resolution stability during automated volume acquisition.

### Reduced milling voltage affects volume-imaging throughput

To determine the effect of reduced FIB accelerating voltage on volume-imaging throughput we analyzed the ASV run in more detail (Supplementary Table 3). After accounting for SEM acquisition time and milling pattern size, normalized non-imaging time increased at reduced voltage in both biological replicates for xenon (45% and 75%), argon (56% and 64%) and nitrogen (28% and 50%) (Supplementary Table 4). In contrast, oxygen showed the opposite trend, with normalized non-imaging time decreasing by 18% and 32% at 15 kV. These results indicated that the effect of reduced voltage-milling on ASV acquisition cycle was ion-species dependent.

## Discussion

Our results establish that cryo-pFIB-SEM block-face quality is determined by the coupled effects of ion species, accelerating voltage, and local specimen composition. Lowering the milling voltage reduces predicted ion penetration and displacement damage^14^ and suppresses charging and voltage-dependent topographic artifacts, but the extent to which voltage can be reduced is constrained by species-dependent curtaining. Conversely, oxygen and nitrogen produced greater membrane sharpness and organelle contrast but were more susceptible to curtaining as accelerating voltage was reduced. These competing effects preclude a single milling condition that is optimal for all specimens and imaging objectives. Instead, the consequences of a given milling regime depend on both the ion beam and the composition and ultrastructure of the biological material being exposed.

The implications of accelerating voltage in cryo-pFIB-SEM volume imaging differ from those in cryo-FIB preparation of lamellae for cryo-ET. In lamella preparation, subsurface ion damage is critical because the remaining specimen is subsequently interrogated by transmission electron microscopy^12,13,15–17^. In volume imaging, by contrast, each milling step generates a new surface that is immediately imaged by SEM. The morphology, composition, charging behavior, and surface potential of this outermost block-face therefore become direct determinants of the information recorded in each image^4,8,18,19^. Our simulations predicted substantially reduced ion penetration and atomic displacement at the species-specific low-kV conditions, while FRC analysis revealed no corresponding loss of SEM resolution. Mean and spatially distributed patch-FRC measurements were broadly similar between high- and low-kV conditions. During automated volume acquisition, however, their temporal behavior differed as FRC-derived resolution deteriorated modestly at 30 kV (∼1 nm over 100 images) but remained stable under low-kV conditions. Although individual runs varied and the underlying mechanism remains unresolved, these results indicate that reducing milling voltage can alter the state of the block-face without compromising SEM spatial resolution and may improve resolution stability during extended acquisition.

The more substantial effects of milling conditions were therefore observed not in nominal spatial resolution but in local image properties, including sharpness, contrast, charging, and surface morphology. Oxygen and nitrogen consistently produced higher membrane sharpness than xenon and argon, while nitrogen produced particularly high contrast in late endosomes. Off-axis imaging provides a possible structural basis for some of these differences: compositional interfaces exhibited species-dependent surface texturing consistent with differential sputtering^14,20^, while reactive ion species may additionally interact with the exposed biological surface^21,22^. Because block-face chemistry was not directly assessed, chemical modification cannot presently be distinguished from differential physical sputtering or associated changes in secondary-electron (SE) yield. These possibilities therefore remain potential interpretations rather than demonstrated sources of the observed contrast.

Importantly, increased sharpness or contrast should not automatically be equated with greater structural fidelity. At 30 kV, bright-line features occurred repeatedly at compositional interfaces, including the GIS-cell boundary, carbon support interfaces, lipid droplets and lipid-rich domains within late endosomes. Off-axis imaging showed that at least some of these features coincided with physical indentations in the block-face, demonstrating that the enhanced signal was associated with topographic modification rather than solely with changes in SEM intensity. Their occurrence with all four species and suppression at lower voltage identify beam energy as a major determinant of this effect. Redeposition, differential sputtering, local charging and changes in the SE yield could all contribute^4,9,19,23^, but the present data do not distinguish their relative contributions. Regardless of the precise mechanism, these observations demonstrate that high apparent contrast at a compositional boundary can coexist with measurable modification of the underlying block-face.

The consequences of such artifacts were particularly apparent in lipid-rich cellular compartments. Whereas whole-cell measurements showed relatively modest voltage-dependent differences, late endosomes displayed stronger changes in charging and contrast. Low-kV milling, particularly with oxygen and nitrogen, reduced charging and restored membrane boundaries that were poorly defined or obscured at 30 kV. This spatial specificity is important because averaging image-quality measurements over an entire cell can conceal substantial local degradation of biological information. Lipid-rich organelles, multilamellar structures, and other compositionally heterogeneous compartments may therefore provide particularly informative tests of milling conditions for cryogenic volume imaging^24^.

Curtaining provides the principal constraint on voltage reduction^25^. Xenon and argon maintained sufficiently uniform block-faces to approximately 10 kV, whereas oxygen and nitrogen required approximately 15 kV under the conditions tested here. Thus, lowering accelerating voltage cannot be considered independently of ion species. Reactive species provide advantages in membrane sharpness and organelle contrast but reach their practical low-voltage limit earlier, whereas xenon and argon permit operation at lower accelerating voltages with less severe curtaining. These relationships define distinct operating regimes rather than a universal hierarchy among ion species.

The practical consequences of these tradeoffs became particularly clear in the 3D datasets. ML-based segmentation showed that milling conditions influenced which organelle features could be recognized in volume data. Noble-gas datasets gained intraluminal detail at low-kV milling, while oxygen and nitrogen provided greater structural definition overall, with low-kV milling reducing charging while retaining fine ultrastructure. These observations connect the 2D image-quality measurements directly to biological information recovery as milling conditions influence not simply how a block-face appears, but which ultrastructural features can ultimately be recognized and analyzed in 3D.

This distinction also highlights a limitation of using nominal spatial resolution alone to characterize cryo-pFIB-SEM dataset quality. FRC-derived resolution changed little between high- and low-kV conditions, whereas organelle-level measurements and ML-based segmentation revealed substantial differences in feature resolvability. Thus, the biological information content of a volume can change without a corresponding change in nominal resolution. Charging, local contrast, membrane definition, and milling-induced surface morphology may instead determine whether a structure can be distinguished from its surroundings and therefore accessed by subsequent segmentation and quantitative analysis.

Accelerating voltage also affected the practical throughput of automated volume acquisition. Non-imaging time increased at reduced voltage for xenon, argon and nitrogen in both biological replicates, with the greatest increase observed for xenon. This effect persisted despite comparable or, for some species, slightly higher beam currents at reduced voltage, as determined by the pFIB column optics for the same aperture index. Oxygen showed the opposite trend, demonstrating that the effect of accelerating voltage on acquisition throughput is species dependent. Because the non-imaging interval includes FIB milling together with other ASV operations, it cannot be interpreted as a direct measure of milling rate. Nevertheless, these differences identify acquisition throughput as an additional consideration when selecting milling voltage for automated volume imaging.

Together, these findings have direct implications for practical cryo-pFIB-SEM operation (Fig. 6). Xenon and argon maintain relatively uniform milling over a broader range of accelerating voltages, whereas oxygen and nitrogen provide enhanced membrane definition and organelle contrast but become increasingly susceptible to curtaining as accelerating voltage is reduced. The choice of ion species and accelerating voltage therefore requires balancing block-face quality and ultrastructural information recovery against milling behavior and acquisition throughput. Reactive species may be preferable when membrane definition and organelle contrast are the primary considerations, whereas the higher sputtering efficiency of xenon and argon^10^ may be advantageous when rapid material removal and imaging larger specimen volumes are priorities.

**Figure 6.**
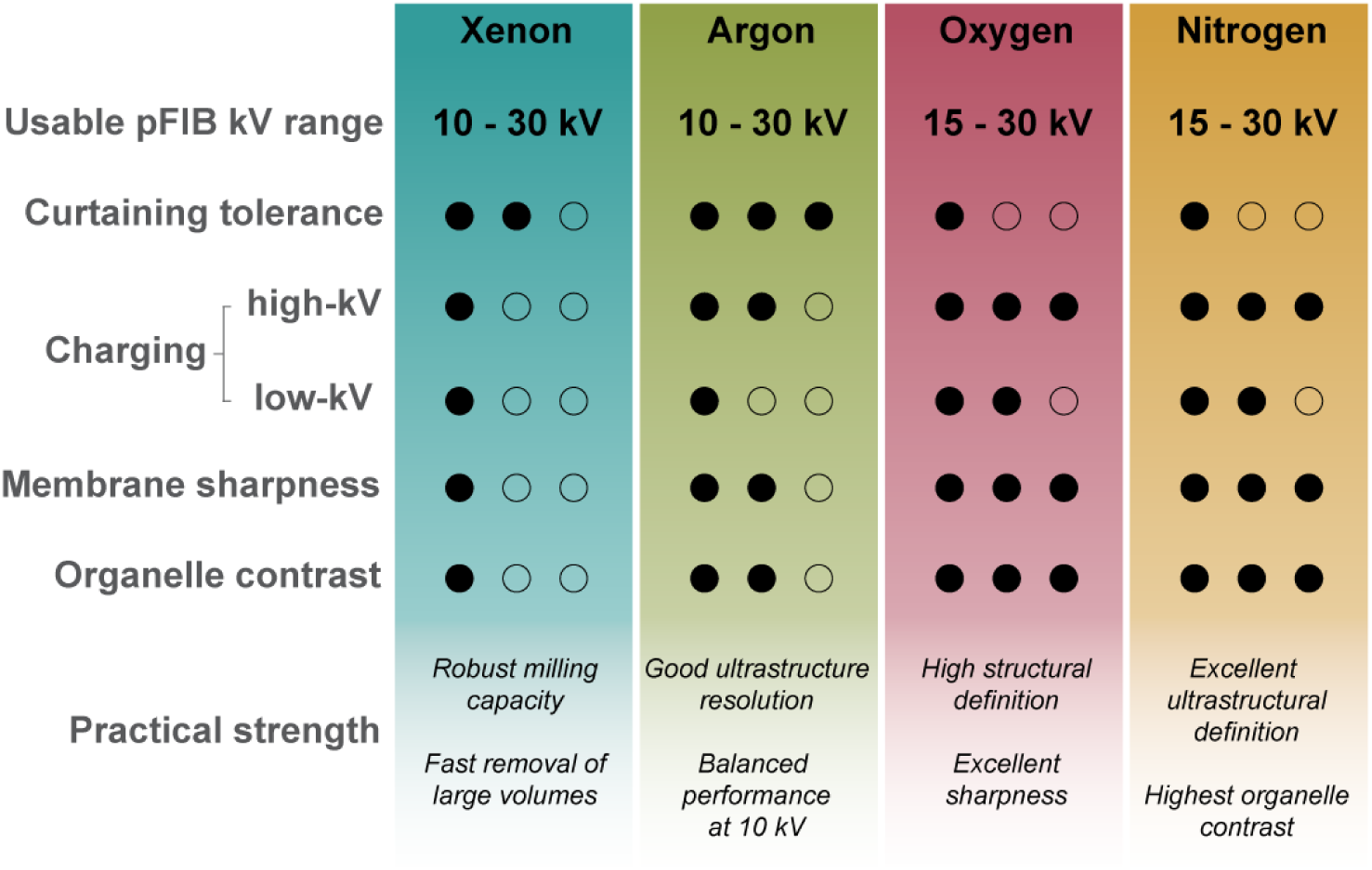
Comparative performance of plasma ion species for cryo-pFIB milling. Qualitative summary of milling and imaging performance for the four plasma ion species integrating the results of curtaining and block-face image-quality analyses. Columns represent individual ion species, and rows summarize key performance criteria. The usable pFIB kV range denotes the accelerating-voltage window over which each ion species yielded acceptable milling quality, bounded by the lowest voltage at which curtaining remained tolerable and by the standard upper milling condition of 30 kV. For curtaining tolerance, membrane sharpness, and organelle contrast, filled and open circles denote more and less favorable performance, respectively, on a three-point relative scale. For charging, however, the scale is inverted: a greater number of filled circles indicates more severe charging rather than better performance. Xenon and argon exhibited minimal charging at both voltages, whereas oxygen and nitrogen showed pronounced charging at 30 kV that was substantially reduced at lower kV. Practical strength summarizes the principal qualitative advantage of each ion species based on the quantitative comparisons presented above.

Under the conditions examined here, argon at 10 kV represents a useful intermediate regime, combining improved ultrastructural definition with a sufficiently uniform block-face while avoiding the stronger low-voltage curtaining observed with oxygen and nitrogen. This should be regarded as a practical operating point rather than a universal optimum, however, because the balance among these effects is likely to depend on specimen composition, beam current, slice thickness, milling geometry, and acquisition duration.

Several variables could further shift or expand these operating regimes. Beam current and slice thickness are likely to influence both milling efficiency and the accumulation of surface artifacts, while milling geometry and rocking or multi-angle strategies may mitigate curtaining and extend the usable low-voltage range^26,27^. SEM accelerating voltage represents an additional independent variable that may interact with the surface state generated by different milling conditions^19^. Direct characterization of block-face chemistry and topography will also be important for distinguishing among differential sputtering, chemical modification, charging and SE effects underlying the species-dependent contrast and surface morphology observed here.

Together, our findings challenge the routine use of 30 kV as a universal default for cryo-pFIB-SEM volume imaging. Rather than identifying a single optimal ion species or accelerating voltage, the results define operating regimes that balance material-removal efficiency against curtaining, charging, surface fidelity and ultrastructural contrast. Lowering accelerating voltage can suppress charging and voltage-dependent topographic artifacts without compromising SEM spatial resolution and may improve resolution stability during prolonged acquisition, while ion species determines both the attainable low-voltage range and the contrast characteristics of the resulting block-face. Matching these operating regimes to specimen composition, volume requirements and the biological target being sought should improve both the efficiency and information content of cryo-pFIB-SEM volume imaging.

## Data availability

The code is available at https://github.com/stjude-cbi/cryoinspect-metrics and the raw EM data are currently available upon request.

## Acknowledgements

The authors would like to thank Scott Blanchard and Anand Singh from the St. Jude Children’s Research Hospital for providing NMuMG cells.

## Author contributions

S.S. designed research with input from C.T., R.K., A.K. and G.S.; C.T. acquired cryo-pFIB-SEM data; C.B., K.V., S.S. and Ali K. worked on image analysis; C.B. and J.H. contributed segmentation and visualization of the vEM datasets; M.Z.Q. prepared vitrified samples; G.S. supervised the project; S.S. and G.S. wrote the paper with contributions from all authors.

## Methods

### Cell culture

Namru Mus musculus mammary gland (NMuMG) cells were cultured in Dulbecco’s Modified Eagle Medium (DMEM, Gibco/Invitrogen), 10% fetal bovine serum (FBS, Gibco/Invitrogen), GlutaMAX (Gibco/Invitrogen), penicillin–streptomycin (PenStrep, Gibco/Invitrogen) and insulin 10 µg/mL (Sigma-Aldrich, USA).

### Sample preparation

Electron microscopy (EM) grids (Quantifoil R 1.2/1.3, Au 200) were glow discharged using a PELCO® easiGlow™ Glow Discharge Cleaning System (Ted Pella, Inc., Redding, CA, USA) and sterilized by incubation in 70% (v/v) ethanol (Sigma-Aldrich, St. Louis, MO, USA) for 5 min, followed by multiple rinses with ultrapure water (Millipore Sigma, Burlington, MA, USA) to remove residual ethanol. Alternatively, grids were sterilized by exposure to ultraviolet (UV) light in a biosafety cabinet for 30 min.

For extracellular matrix coating, grids were placed on droplets of 10 μg/ml fibronectin (Sigma-Aldrich, USA) in phosphate-buffered saline (PBS, Thermo Fisher Scientific, Waltham, MA, USA) inside an Ibidi 35 mm glass bottom dish (Ibidi GmbH, Gräfelfing, Germany). Additional 5 μl of fibronectin solution was applied to the top of each grid. Grids were incubated for 30 min at 37 °C and gently rinsed three times with PBS.

Approximately 5 μl of suspension of NMuMG cells was applied to each coated grid. Cell concentration was adjusted to achieve a seeding density of approximately one to two cells per grid square and assessed by light microscopy after incubation for 30 min. After cell seeding, grids were immersed in 2 ml of conditioned (2-3 days old) culture medium and incubated overnight at 37 °C in a humidified atmosphere containing 5 % CO₂.

Grids were vitrified using a Leica EM GP2 Automatic Plunge Freezer (Leica Microsystems, Vienna, Austria). Individual grids were mounted in the plunge freezer and back-blotted under controlled environmental conditions (25 °C and 60% relative humidity) before being rapidly plunged into liquid ethane. Vitrified grids were transferred to liquid nitrogen for storage.

### Cryo-pFIB-SEM

All experiments were performed using a Thermo Scientific Hydra Bio Plasma-FIB in UX configuration, focused ion beam-scanning electron microscope equipped with a multi-species plasma ion source (Xe, Ar, O and N), an electron column with a magnetic immersion lens, a platinum (Pt) gas injection system (GIS) operated at 28 °C, and a Pt micro-sputter coater. Vitrified grids were clipped into TFS AutoGrids under cryogenic conditions, loaded into a 35° pre-tilted AutoGrid shuttle and transferred to the microscope. The cryo-stage and shield were maintained at approximately −185 °C and −192 °C, respectively.

SEM tile sets of each grid were acquired using TFS Maps 3.36 at a stage tilt of 35°, at which the electron beam was normal to the grid. Tile sets were used to select regions of interest (ROIs), with each ROI corresponding to a single cell. Block-faces were prepared by manual milling at a stage tilt of 35°, corresponding to a pFIB incidence angle of 38° relative to the grid plane (Fig. 1b). SEM images were subsequently acquired automatically after each milled slice using Auto Slice and View (ASV) 5.13. Unless otherwise stated, images were acquired at a voxel size of 5 × 5 × 20 nm, with the SEM operated in immersion mode at a landing energy of 1.2 keV and beam current of 13 pA. Images were collected using the through-lens detector (TLD) in down-hole-voltage (DHV) mode (−245 V suction tube, −15 V mirror), with a dwell time of 50 ns, line integration of 150 and frame integration of 1. Detector contrast and brightness settings were held constant across comparative imaging experiments.

### pFIB low-voltage screening

To determine the usable low-kV milling range for each ion species, grids were coated with approximately 600 nm of Pt deposited by GIS, followed by a thin micro-sputtered Pt layer (120 nA Xe beam, 120 s). For each ion species, cross-sections were milled at three independent cellular ROIs across seven accelerating voltages, 30, 20, 15, 10, 8, 5 and 2 kV, with ASV used for automated SEM acquisition (Supplementary Table 1). For each species, experiment started with the highest voltage value and the same aperture index was used throughout the voltage series.

### Feature-to-feature comparison

For paired feature-to-feature comparisons of 30-kV and low-kV milling (Fig. 2a), grids were coated with alternating layers of GIS-deposited and micro-sputtered Pt in the following sequence: 700 nm GIS Pt, micro-sputtered Pt, 150 nm GIS Pt, micro-sputtered Pt, 150 nm GIS Pt and micro-sputtered Pt. Each micro-sputtered layer was approximately 10-20 nm thick (120 nA Xe beam, 120 s). This multilayer coating enabled use of the ASV dynamic autofunctions when required.

Five cellular ROIs were analyzed per ion species. At each ROI, up to five regions distributed through the cell volume were imaged, with each comparison comprising at least five SEM images acquired at each of two pFIB accelerating voltages: 30 kV and the corresponding species-specific low-kV condition (10 kV for Xe and Ar; 15 kV for O and N). Milling patterns were positioned such that the final slice acquired at one voltage was immediately followed by the first slice acquired at the second voltage, thereby maintaining near-continuous sampling through the organelle of interest. SEM images were acquired automatically using ASV.

### Block-face damage assessment

Following tile-set acquisition, grids were coated with approximately 600 nm of GIS-deposited Pt followed by a thin micro-sputtered Pt layer (120 nA Xe beam, 120 s). Stage tilt was maintained at 35° throughout the experiment, with only stage rotation varied. Initial cross-sections were prepared manually by Ar milling at 30 kV and 2 nA using an Si-cleaning cross-section (Si-ccs) pattern with a depth of 4 µm. After changing to the experimental milling condition, approximately 400 nm of block-face material was removed before SEM imaging.

An on-axis (front-view) secondary-electron (SE) image was acquired at a stage rotation of −70°. The stage was then rotated to 5° to acquire an off-axis (side-view) SE image before being returned to −70° for acquisition of an on-axis backscattered-electron (BSE) image. Although a 35° pre-tilted AutoGrid shuttle was used, the holder was defined as “Standard Shuttle” in the xT Microscope Control software to permit the stage rotation required for off-axis imaging. Images were acquired at a beam current of 13 pA without frame integration; additional SEM conditions are provided in Supplementary Table 2.

### Volume imaging

Following tile-set acquisition, grids were coated with alternating GIS-deposited and micro-sputtered Pt layers in the following sequence: 450 nm GIS Pt, micro-sputtered Pt, 450 nm GIS Pt and micro-sputtered Pt. Each micro-sputtered layer was approximately 10-20 nm thick (120 nA Xe beam, 120 s). Two independent cellular ROIs were selected per ion species. At each ROI, approximately 100 consecutive SEM images were acquired at each of two milling voltages: 30 kV and the corresponding species-specific low-kV condition (10 kV for Xe and Ar; 15 kV for O and N). Initial cross-sections were prepared manually, after which volumes were acquired automatically using ASV.

Acquisition timing was determined from metadata embedded in the SEM images acquired during automated slice-and-view (ASV) volume imaging. To standardize the analysis across datasets, 100 slices of each volume were considered. The interval between acquisition timestamps of consecutive SEM images was calculated. The SEM frame acquisition time was subtracted from each remaining interval to estimate the non-imaging component of the ASV cycle, which includes FIB milling and other operations occurring between consecutive SEM acquisitions. Mean non-imaging time was calculated for each dataset (Supplementary Tables 3 and 4). Because the lateral extent of the milled region varied between datasets, mean non-imaging time was normalized to the corresponding milling length and expressed as s µm⁻¹. Comparisons between accelerating voltages were made within biological replicates.

### FRC calibration data collection

For calibration of Fourier ring correlation (FRC) measurements, the cryo-stage was maintained at approximately 0 °C to improve chamber vacuum, and a TFS high-resolution gold-on-carbon standard was loaded using a standard flat shuttle. Images were acquired at a working distance of 3.5 mm and stage tilt of 0°. SEM acquisition parameters were immersion mode, TLD detector in DHV mode, 1.2 keV landing energy, 13-pA beam current, 10 µs dwell time, line integration of 1 and an image size of 2100 × 2100 pixels.

Images were acquired at pixel sizes of 5, 2.5 and 1 nm, corresponding to horizontal field widths of 10.5, 5.25 and 2.1 µm, respectively. Five independent ROIs were selected at each pixel size. Different ROIs were used at each pixel size to avoid repeated electron exposure and associated contamination, giving 15 ROIs and 30 images in total. At each ROI, focus and astigmatism were manually optimized before two consecutive images were acquired without further adjustment.

### Image analysis

#### Curtaining

Curtaining was quantified by isolating its characteristic directional signal in the frequency domain, following Fourier-based approach for detecting stripe artifacts in FIB-SEM images^28,29^. Vertical intensity bands produce enhanced spectral power near the horizontal frequency axis. Curtain-associated frequencies were therefore defined using a symmetric wedge extending ±5° from this axis. Very low spatial frequencies were excluded using a normalized radial-frequency lower limit of 0.02, while an upper limit of 1.0 retained the remaining sampled frequencies. Horizontal and vertical frequency coordinates were normalized by image width and height before calculating the wedge angle, preventing image dimensions and aspect ratio from distorting the wedge orientation.

The global curtaining score was calculated as the percentage of spectral energy within the analyzed frequency range that occurred inside the directional wedge:

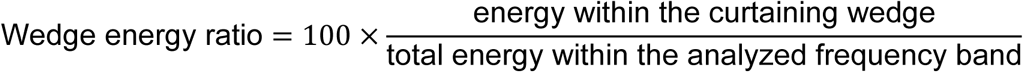

Higher values indicate a greater contribution from vertically aligned intensity variation. Frequencies within the wedge were inverse Fourier transformed to reconstruct the directionally aligned image component.

The spatial distribution of curtaining was estimated from the local root-mean-square (RMS) amplitude of this reconstructed component, calculated within a 15 × 15-pixel neighborhood. Local image texture was estimated by subtracting a 63 × 63-pixel boxcar-smoothed image from the original and calculating the RMS amplitude of the residual within the same neighborhood. The local curtain-to-texture amplitude ratio was calculated as:

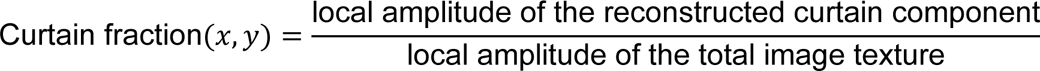

Pixels with a ratio greater than 0.35 were classified as curtain-affected regions. Candidate thresholds of 0.25, 0.35, and 0.50 were visually evaluated. A threshold of 0.35 was selected as the most robust compromise, minimizing noise-related detections while retaining visibly curtain-affected regions.

### Charging and sharpness

Whole-cell and organelle masks were generated using a custom plugin for napari^30^. For charging and sharpness measurements, pixels within those masks were classified as foreground, background or charging-associated using ilastik (v1.4.2)^31^. For every image of every gas and voltage the pixel classification was performed and probability and segmentation map generated for each, from which we extracted the needed pixel labels that were used to quantify sharpness and the percent of charging pixel inside the cell or organelle.

To test the robustness of our sharpness measurements with respect to uncertainty in pixel classification, we calculated sharpness using foreground and background pixels selected at probability thresholds of 0.25, 0.5, and 0.75. Varying the threshold changes the confidence required for a pixel to be assigned to the respective class, with higher thresholds restricting the analysis to more confidently classified pixels. This comparison allowed us to evaluate whether the measured sharpness was sensitive to the choice of classification threshold. A threshold of 0.50 was used for subsequent quantitative analysis.

Image sharpness was quantified using the Tenengrad focus measure, defined as the mean squared image-gradient magnitude^32–34^. Horizontal and vertical gradients, *G_x_* and *G_y_*, were calculated using 3 × 3 Sobel operators, and sharpness within a region *Ω* was calculated as:

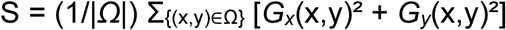

Higher scores indicate stronger local intensity transitions. Because gradient energy is dependent on image content, measurements were restricted to the corresponding cellular or organelle masks; regions outside the cell were excluded.

Charging was quantified using a custom Python script as the fraction of pixels within the corresponding cell or organelle mask classified as charging-associated and is reported as a percentage.

For feature-to-feature comparisons, technical measurements obtained within each cell were first averaged to generate one value per biological replicate for each condition. Paired high- and low-kV measurements from the five biological replicates were then compared using a two-sided Wilcoxon signed-rank test (n = 5 biological replicates). This procedure was applied separately to charging and sharpness measurements for the whole cell and the indicated organelle classes. Heatmaps in Figure 3 display the median of the five biological-replicate means.

### Contrast

To measure contrast, which is created by differences in intensity, we used the robust root mean square metric (RMS)^29,35^. RMS is the standard deviation of the grey values and is reported in raw standard deviations and the coefficient of variation.

The RMS intensity variation was calculated as the standard deviation of gray values where *N* is the number of pixels measured (whole image or inside the cell mask) and *I_i_* is the intensity of pixel *i*. To account for differences in mean image intensity, we also calculated the coefficient of variation, which is dimensionless and therefore facilitates comparison across images with different overall intensity levels.

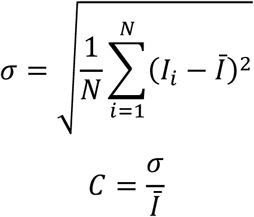

Where

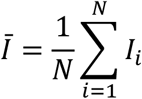

As already described, for feature-to-feature comparisons, technical measurements obtained within each cell were first averaged to generate one value per biological replicate for each condition and paired high- and low-kV measurements from the five biological replicates were then compared using a two-sided Wilcoxon signed-rank test (n = 5 biological replicates). Heatmaps in Figure 3 display the median of the five biological-replicate means.

### FRC measurement

Image resolution was estimated using a checkerboard-based single-image FRC approach, with single-image estimates calibrated against conventional two-image FRC measurements obtained from independently acquired repeat images of the same field of view. For single-image FRC, each SEM image was divided into two interleaved sub-images using a checkerboard split^9,36^ and FRC was calculated between the resulting sub-images over concentric spatial-frequency rings. Resolution was defined as the first persistent crossing below the fixed 0.143 threshold^37,38^, with the crossing frequency determined by linear interpolation. The corresponding spatial period was converted to the original-image pixel scale and subsequently corrected using the calibration described below.

Following the calibration strategy described by Dumoux et al.^9^, calibration was performed using 15 independently acquired image pairs, comprising five fields of view at each of three pixel sizes (1, 2.5 and 5 nm per pixel). Repeat images were registered by phase cross-correlation. Reference resolution (*r_ref_*) was obtained by calculating conventional two-image FRC between each registered image pair. Raw single-image resolution (*r_co1_*) was calculated independently for each image using the checkerboard method and averaged within each image pair. Calibration relationships were determined for both complete image crops and non-overlapping 256 × 256-pixel patches. All subsequent specimen measurements used patch-based FRC, corrected using the patch calibration curve.

Specimen images were divided into non-overlapping patches, and only patches intersecting the full-cell mask were included. Standard specimen images were analyzed using 256 × 256-pixel patches. For 1000 × 1000-pixel volume images, 250 × 250-pixel patches were used to permit exact tiling in both dimensions. The 256 × 256-pixel calibration relationship was applied to both patch sizes. Calibrated spatial periods were converted to nanometers, and both raw *r_ref_* and calibrated *r_co1_* estimates were retained.

We compared FRC resolution measurements using 250 × 250 and 256 × 256-pixel patches across 800 images acquired at 15 and 30 kV while milling with oxygen or nitrogen. For each image, the mean calibrated FRC resolution was calculated from all patches. The two patch sizes produced nearly identical results, with an overall difference of only 0.035 nm (0.20%), indicating that they yield effectively equivalent FRC measurements.

### Temporal analysis of FRC-derived resolution during volume acquisition

To assess the stability of SEM spatial resolution during automated volume acquisition, patch-based FRC-derived resolution was calculated for each consecutive block-face image as described above. The analysis comprised two biological replicates for each combination of ion species and milling-voltage regime, corresponding to 16 acquisition runs. Voltage regime was defined as 30 kV or the corresponding species-specific low-kV condition (10 kV for Xe and Ar; 15 kV for O and N).

Temporal behavior within individual acquisition runs was visualized by plotting FRC-derived resolution against slice number. For each run, an ordinary least-squares linear regression was fitted independently to provide a descriptive estimate of the temporal slope. These individual fits were used for visualization of run-to-run variability (Supplementary Fig. 9) and were not used as independent observations for statistical inference. Because images within the same volume represent repeated measurements from a single acquisition, FRC values were analyzed using linear mixed-effects models in Python 3.11.15 with statsmodels 0.15.0^39^. The final model included ion species, voltage regime, their interaction, slice number, and the slice number × voltage interaction as fixed effects. Slice number was treated as a continuous measure of acquisition progress. Acquisition run (ion species × voltage regime × biological replicate) was included as a random intercept to account for correlations among images from the same volume and differences in baseline FRC-derived resolution among runs. A random slope for image number was also evaluated but could not be reliably estimated with two biological replicates per condition and was therefore excluded.

To determine whether temporal changes in FRC-derived resolution differed according to ion species or voltage regime, nested models with and without the corresponding image number interaction terms were compared using likelihood-ratio tests. Models used for fixed-effect comparisons were fitted by maximum likelihood. The retained model was subsequently refitted using restricted maximum likelihood (REML) for parameter estimation. Voltage-specific temporal slopes and 95% confidence intervals (CI) were obtained as linear contrasts of the fitted model coefficients using Wald tests. For presentation in Supplementary Figure 10, slopes were multiplied by 100 and expressed as the model-estimated change in FRC-derived resolution per 100 images. Positive values indicate increasing FRC-derived resolution values and therefore deterioration in spatial resolution during acquisition.

Because the default L-BFGS optimizer converged to a boundary solution, model fitting was cross-checked using BFGS, conjugate gradient (CG), Powell and L-BFGS optimization. Reported estimates were based on the mutually consistent solutions obtained with Powell and CG. Residual diagnostics indicated modest right-skew, and the analysis was repeated using log-transformed FRC-derived resolution values as a robustness check, yielding consistent effect directions and statistical conclusions.

### Segmentation and visualization

Segmentation of structures of interest was performed semi-automatically using Dragonfly 3D world (v. 2025.1, Comet Technologies Canada Inc.). Prior to segmentation, cryo-FIB-SEM images were filtered to reduce noise and enhance structural borders for easier and better ground truth generation. First, a Gaussian smoothing was applied in two dimensions to all slices using a kernel size of 3 and standard deviation of 1.0. The Gaussian filter reduces high-frequency noise by replacing pixel intensities with a Gaussian-weighted average of neighboring pixels while providing comparatively good preservation of edges. Secondly, an unsharp filter was applied in two dimensions to all slices using a kernel size of 3, standard deviation of 1.0 and unsharp factor 1. The unsharp filter enhances local contrast at structural boundaries by subtracting a blurred representation of the image from the original image.

Next, to generate 3D segmentations we used the Dragonfly deep-learning tool with U-net architecture^40^. For each acquired volume a separate U-net model in 2.5D mode using five consecutive slices as input was trained with a depth level of 5 and an initial filter count of 64. For each volume, four to five slices throughout the volume that showed the organelles of interest were manually segmented and put as mixed regions, which assigns patches from these regions randomly to training, validation, and evaluation. An additional slice was put as a monitoring region to provide visual feedback on segmentation performance during training.

The models were trained for 100 epochs with a batch size of 64 using cross-entropy loss function and Adadelta optimization algorithms^41^ with a learning rate of 1.0. The patch sizes were 128 x 128 x 5 voxel and 64 x 64 x 5 with a stride ratio of 0.25, depending on workstation used for training. An augmentation factor of 10 was applied and included horizontal and vertical flipping, as well as a rotation of 180°, zoom of 0.9 and 1.10 and shear of 2. The training was performed on two workstations, each equipped with NVIDIA RTX 5000 Ada GPU with 32 GB of memory and 512 GB and 192 GB of RAM, respectively.

After the training the individual models were applied to the corresponding complete volume datasets. As a final step the automated segmentations were manually cleaned up using the 3D-connected components tool. Manual correction was restricted to removal of pixel islands and to only organelles that had been initially detected by the automated segmentation. Visualization and movies were generated using the snapshot option or movie maker tool from Dragonfly 3D world.

### SRIM simulations in vitreous ice

Ion-induced interactions were simulated using SRIM (Stopping and Range of Ions in Matter, version 2013)^42^. Xe and Ar were simulated at 30 and 10 keV, and O and N at 30 and 15 keV, using amorphous H₂O ice (density, 0.94 g cm⁻³) as the target material. For each condition, the ion-incidence angle was set to 89.5° relative to the target surface to approximate the grazing-incidence geometry used during cryo-pFIB milling, and 10 000 ion trajectories were simulated. Default SRIM displacement and surface-binding energies for the target were used. For each condition, the depth distribution of implanted ions and ion-induced target vacancies was extracted, together with the total number of target vacancies generated per incident ion.

---

**Supplementary Figure 1.**
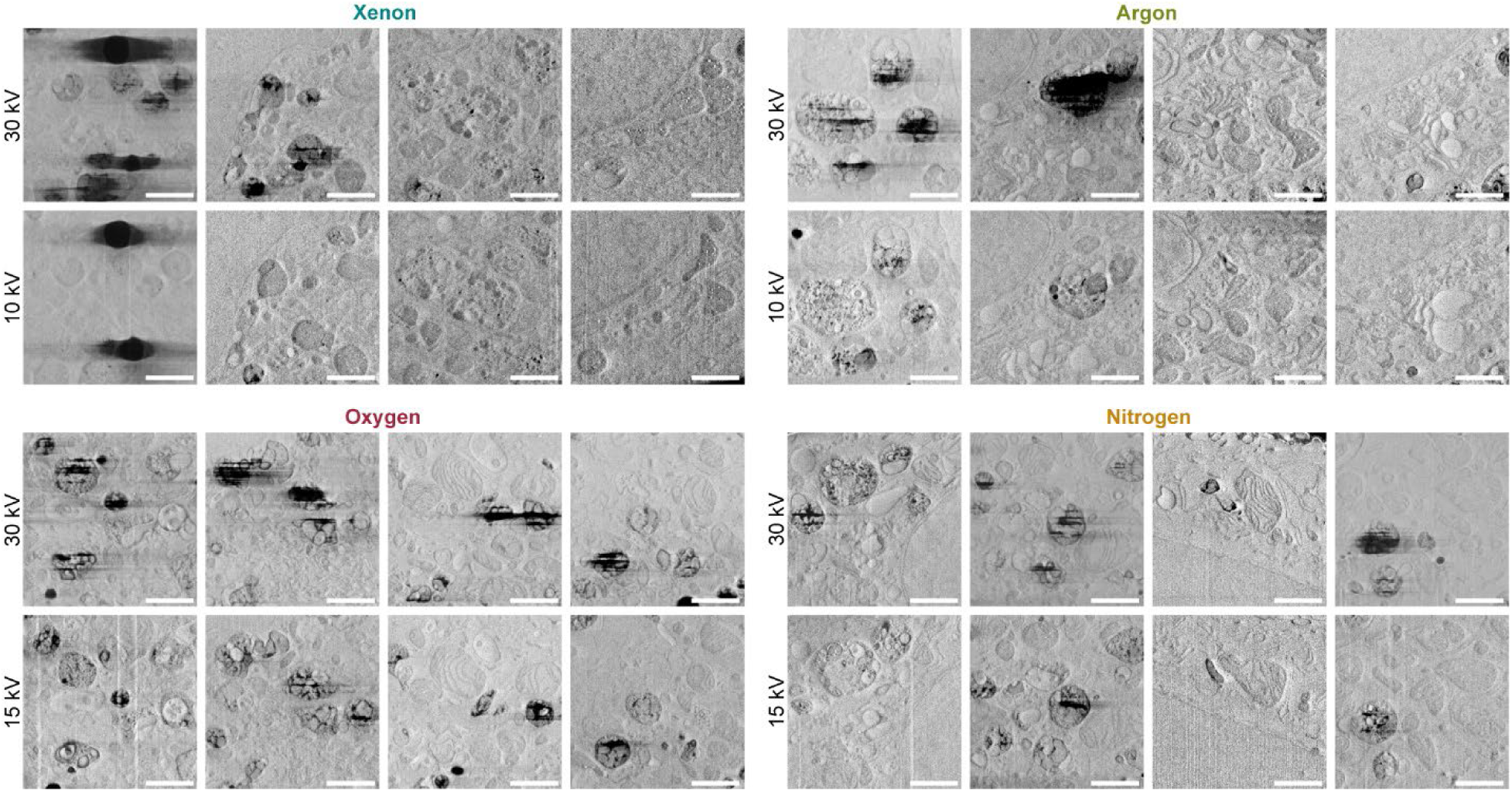
SEM image quality of the same organelles after milling with high- and low-voltage pFIB regimes. Representative SEM micrographs from the ‘feature-to-feature’ comparison workflow described in Figure 2a, showing additional examples across biological replicates. Scale bar: 500 nm.

**Supplementary Figure 2.**
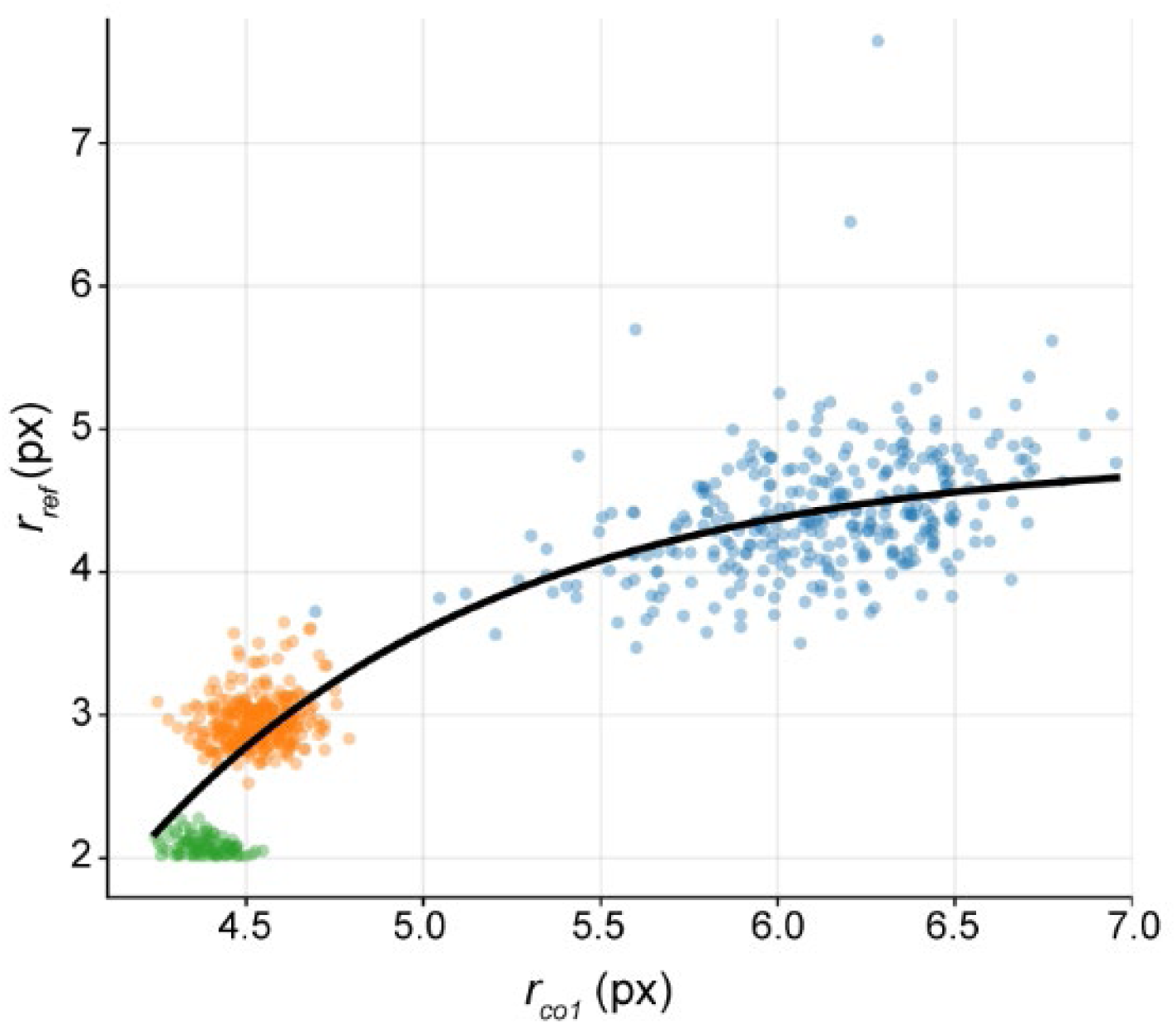
Calibration of single-image Fourier Ring Correlation (FRC)-derived resolution. Calibration curve relating single-image checkerboard FRC-derived resolution to conventional two-image FRC-derived resolution for images acquired on the microscope used in this study. The x-axis shows the resolution estimated from a single image by splitting each 256 × 256-pixel patch into two interleaved checkerboard sub-images and calculating FRC using the 0.143 threshold. The y-axis shows the reference resolution obtained by conventional two-image FRC from independently acquired images of the same field of view. Resolution is expressed as spatial period in pixels; therefore, smaller values indicate higher spatial resolution. Each point represents one 256 × 256-pixel patch and is colored according to the acquisition pixel size (blue, 1.0 nm per pixel; orange, 2.5 nm per pixel; green, 5.0 nm per pixel; n = 727 patches). The fitted relationship was used to calibrate patch-based single-image FRC measurements of experimental datasets.

**Supplementary Figure 3.**
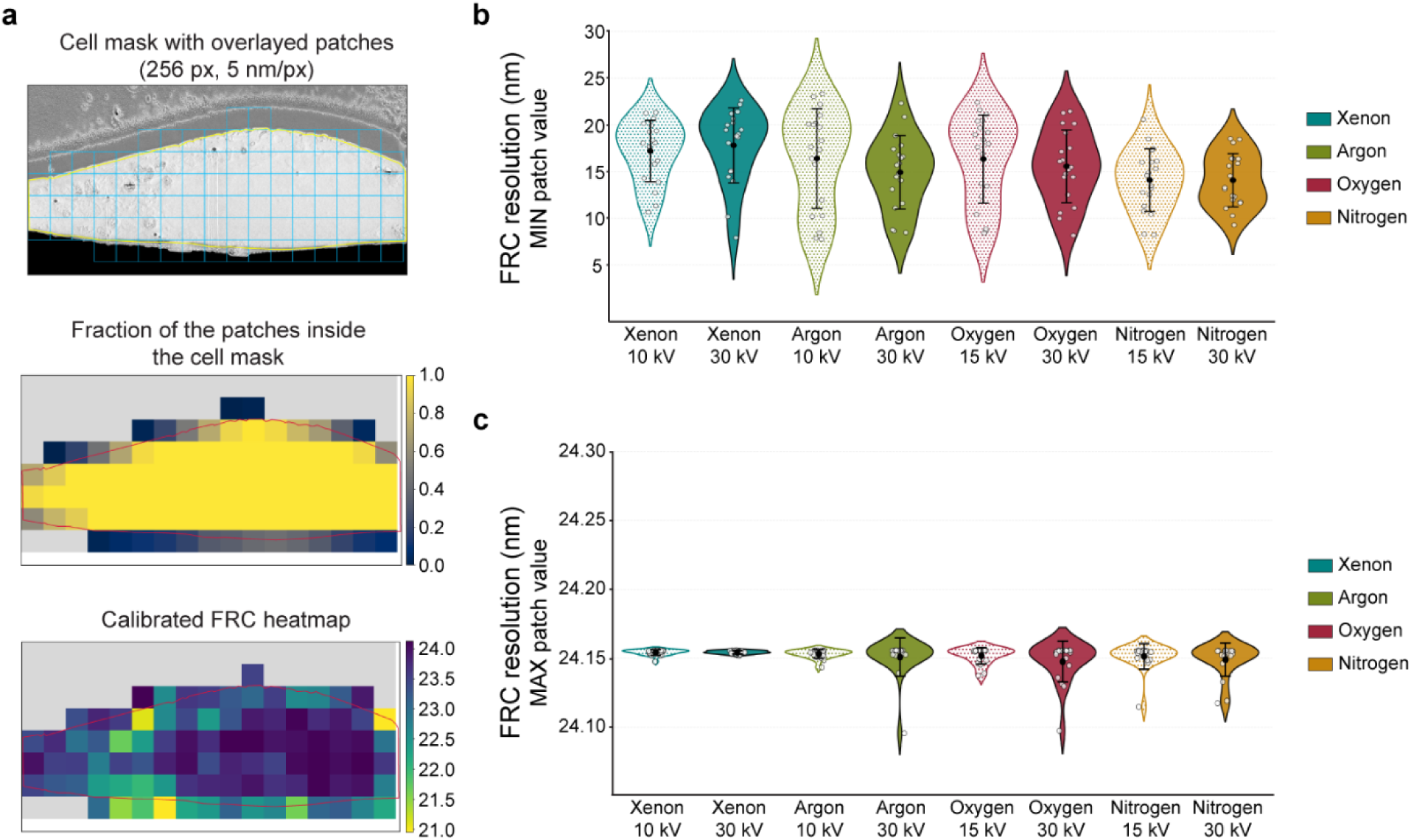
Fourier Ring Correlation (FRC) resolution analysis for SEM micrographs acquired after milling with different ion species operating at high and low milling voltages. (A) Cell mask (yellow outline) was applied to each micrograph and divided into 256 x 256-pixel patches (blue squares) (top). The fraction of each patch inside the cell mask was calculated (middle) and a heatmap with calibrated FRC resolution per each patch generated (bottom). (B) Average highest resolution value per patch. (C) Average lowest resolution value per patch. Per each condition, standard deviation reflects five biological replicates, each with up to 5 technical replicates.

**Supplementary Figure 4.**
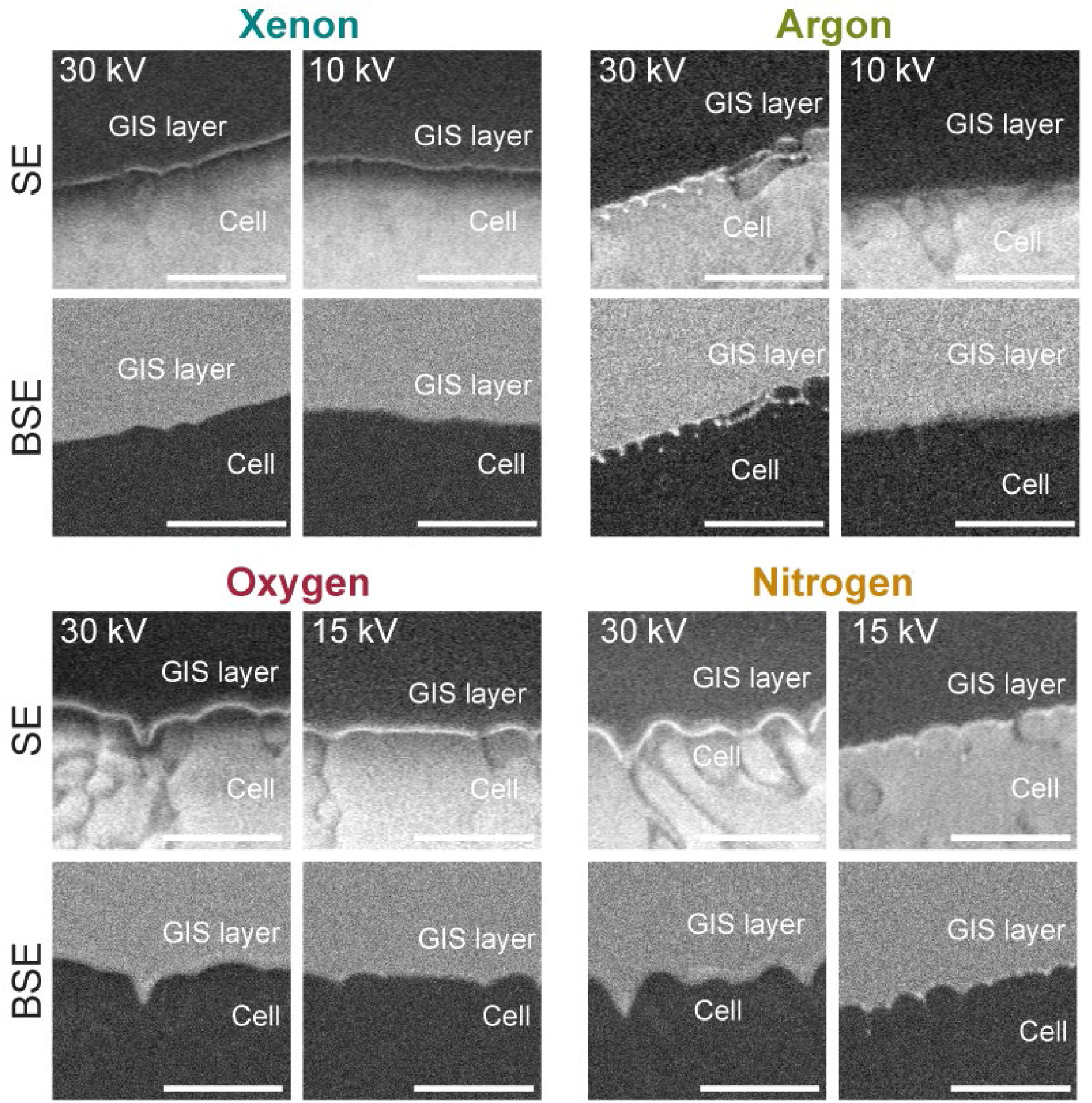
Interface between the cell surface and GIS protective layer. The bright-line feature between the surface of vitrified cell and platinum protective layer, imaged using secondary electron (SE) and backscattered electron (BSE) imaging. Scale bar: 500 nm.

**Supplementary Figure 5.**
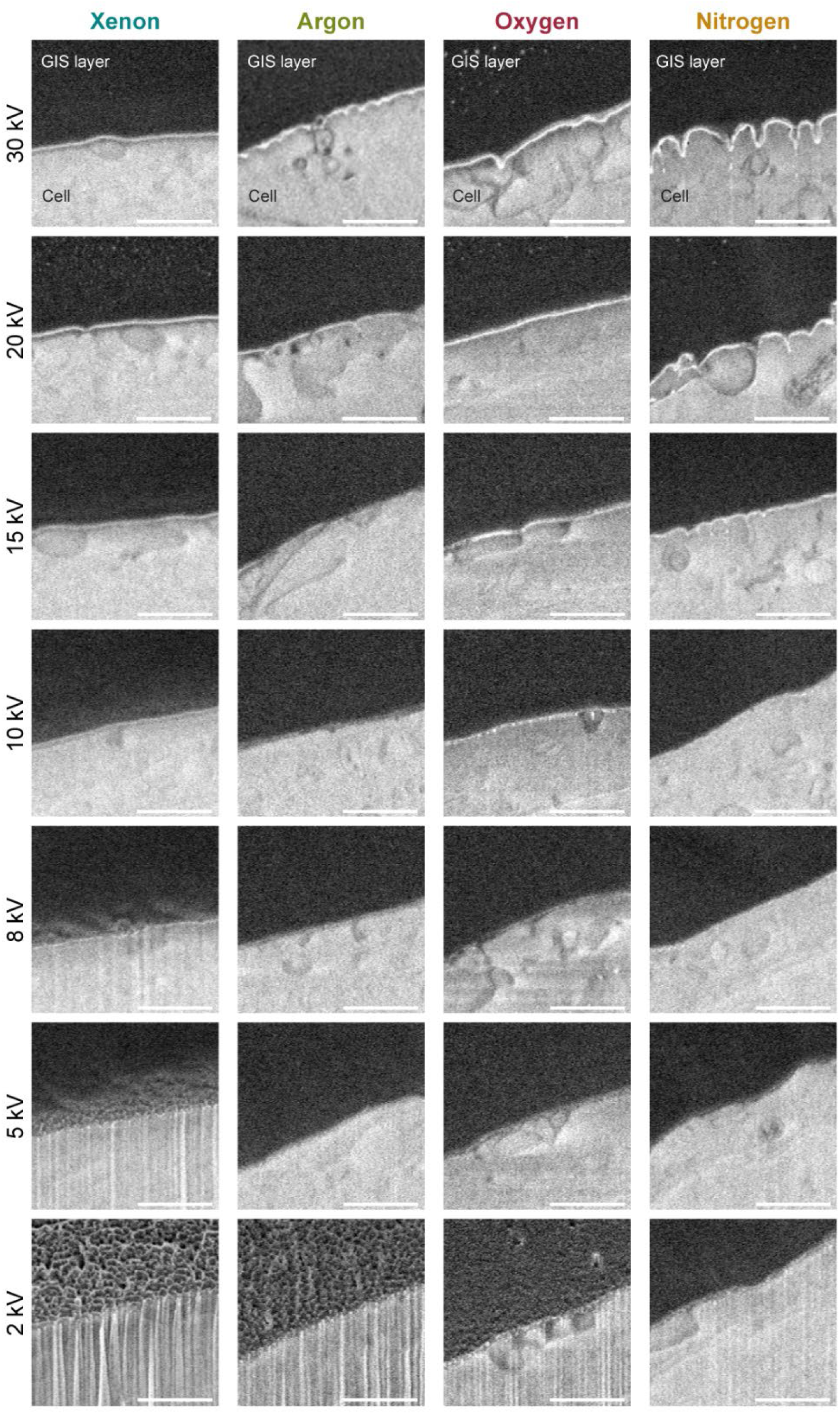
Interface between the cell surface and GIS protective layer. The bright-line feature between the surface of the vitrified cell and platinum protective layer diminishes with low-kV milling. Scale bar: 500 nm.

**Supplementary Figure 6.**
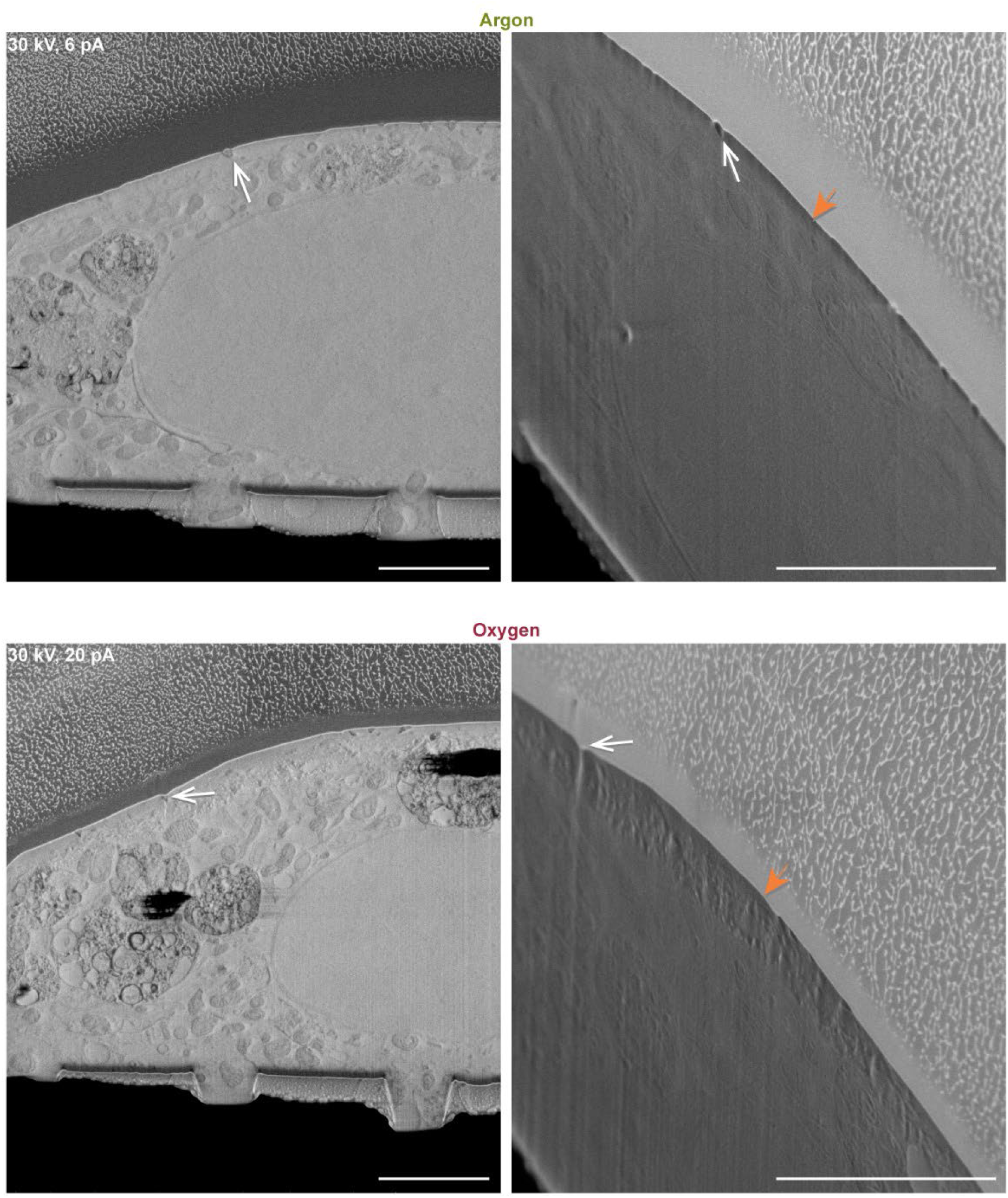
pFIB milling on-axis (left) and off-axis (right) SEM micrographs of the block-face milled with 30 kV and lower ion beam currents compared to the ones used in Figure 4; argon at 6 pA (top) and oxygen at 20 pA (bottom). White arrow indicates the same feature in on-axis and off-axis micrographs and orange arrow indicates undercut under the GIS protective layer. Scale bar: 1 µm.

**Supplementary Figure 7.**
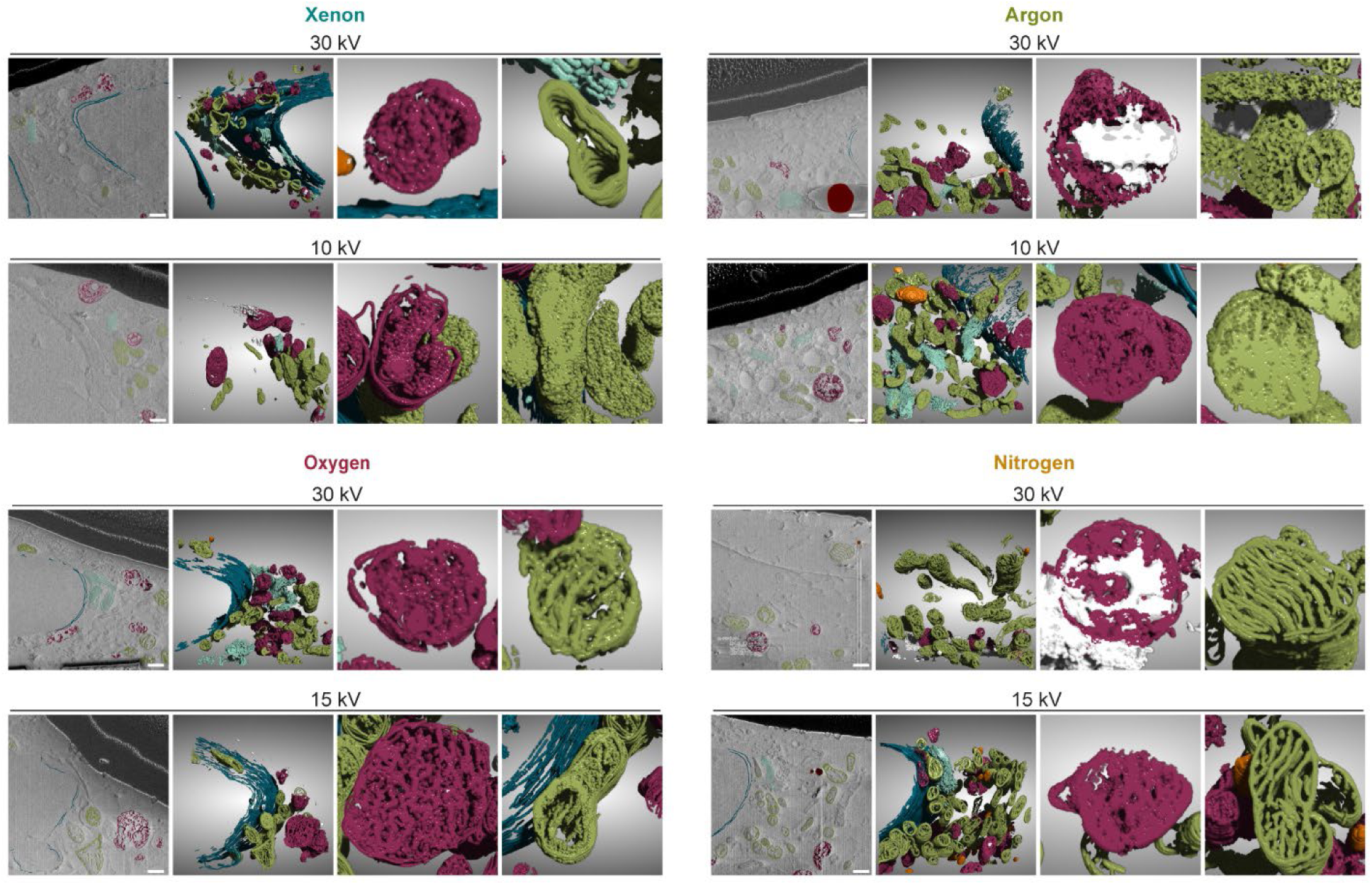
Machine learning-based segmentation and 3D visualization of cellular compartments in the second set of biological replicates. Dark magenta indicates membranes in late endosomes, green indicates membranes associated with mitochondria, dark blue indicates the nuclear membrane, light blue indicates the Golgi apparatus, orange indicates lipid bodies and white indicates charged regions. Scale bar: 500 nm.

**Supplementary Figure 8.**
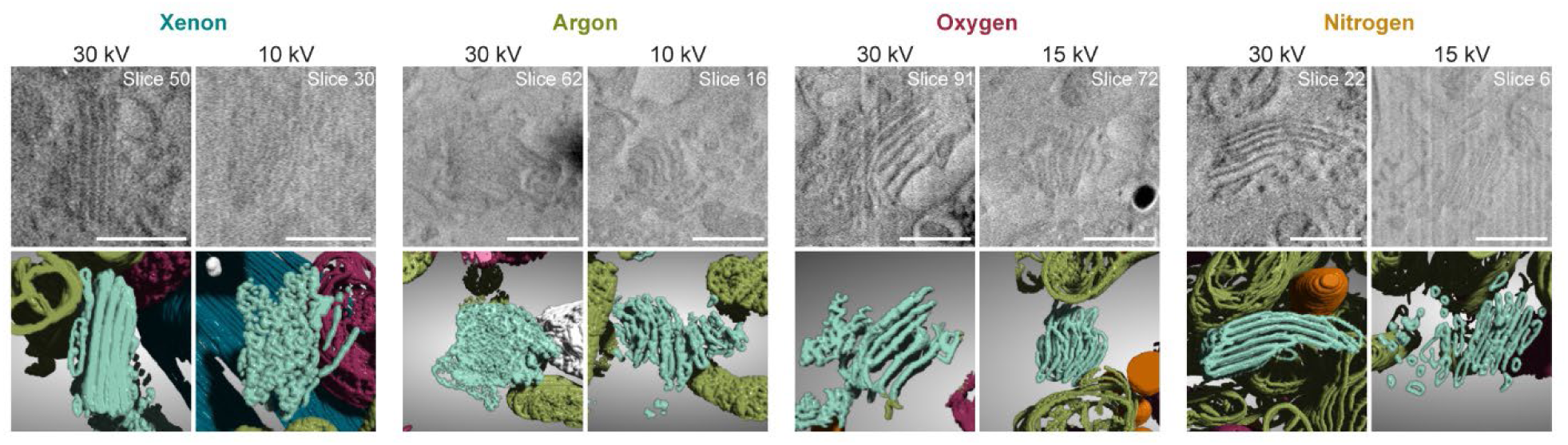
Visualization of the Golgi apparatus. Representative SEM micrographs from volume imaging experiments showing the Golgi apparatus, with corresponding 3D visualization generated by machine learning-based segmentation. Data quality and the resulting structural visualization varied with plasma gas species and milling voltage. Scale bar: 500 nm.

**Supplementary Figure 9.**
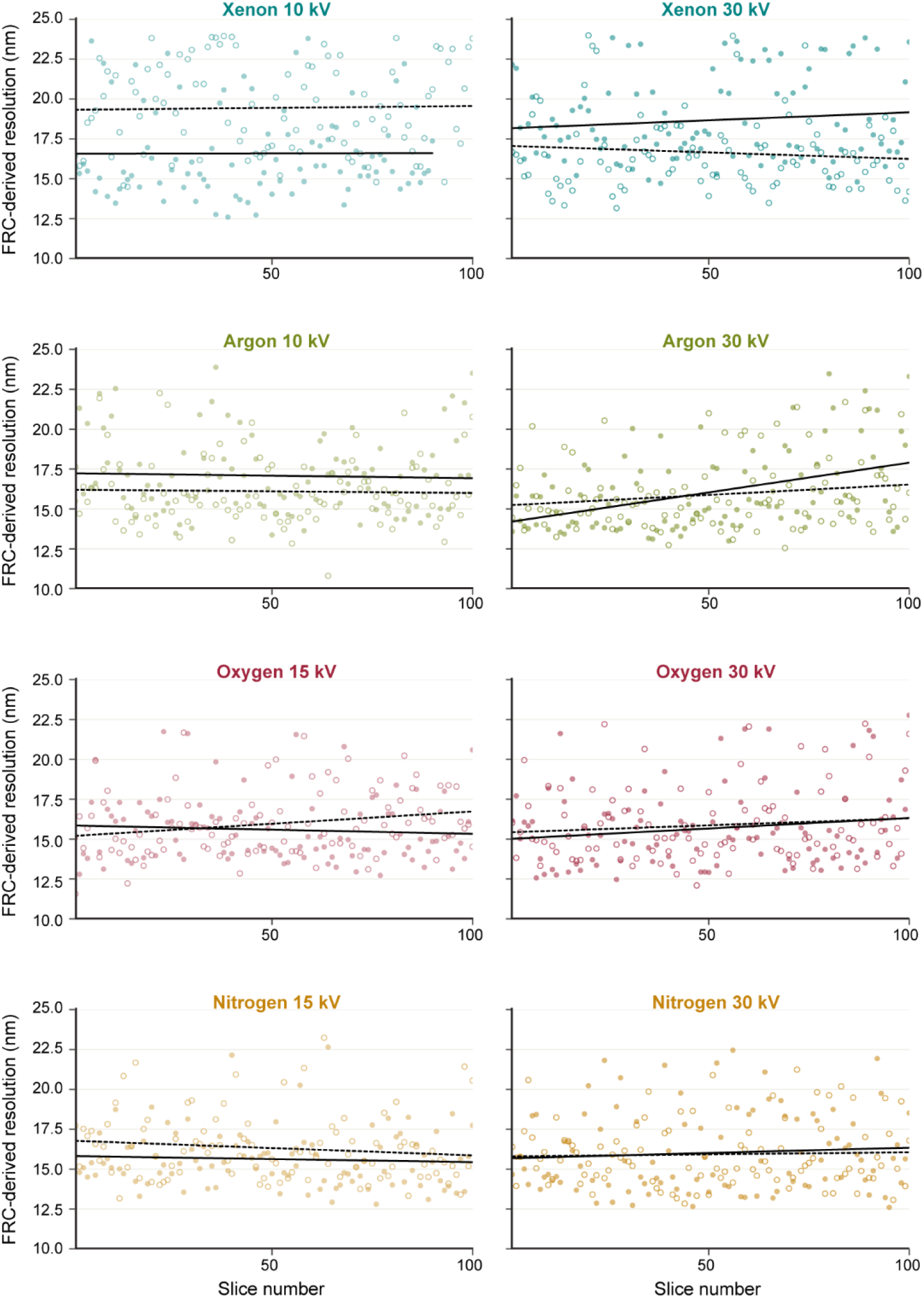
Temporal variation in FRC-derived resolution during automated cryo-pFIB-SEM volume acquisition. FRC-derived resolution was measured for consecutive block-face images acquired using Xe, Ar, O and N at 30 kV and the corresponding species-specific low-kV conditions (10 kV for Xe and Ar; 15 kV for O and N). Filled and open circles represent two independent biological replicates for each condition. Dark grey lines (full – biological replicate 1; dotted – biological replicate 2) show ordinary least-squares fits to individual acquisition runs and are included to visualize run-to-run temporal variation. Lower FRC-derived resolution values indicate higher spatial resolution.

**Supplementary Figure 10.**
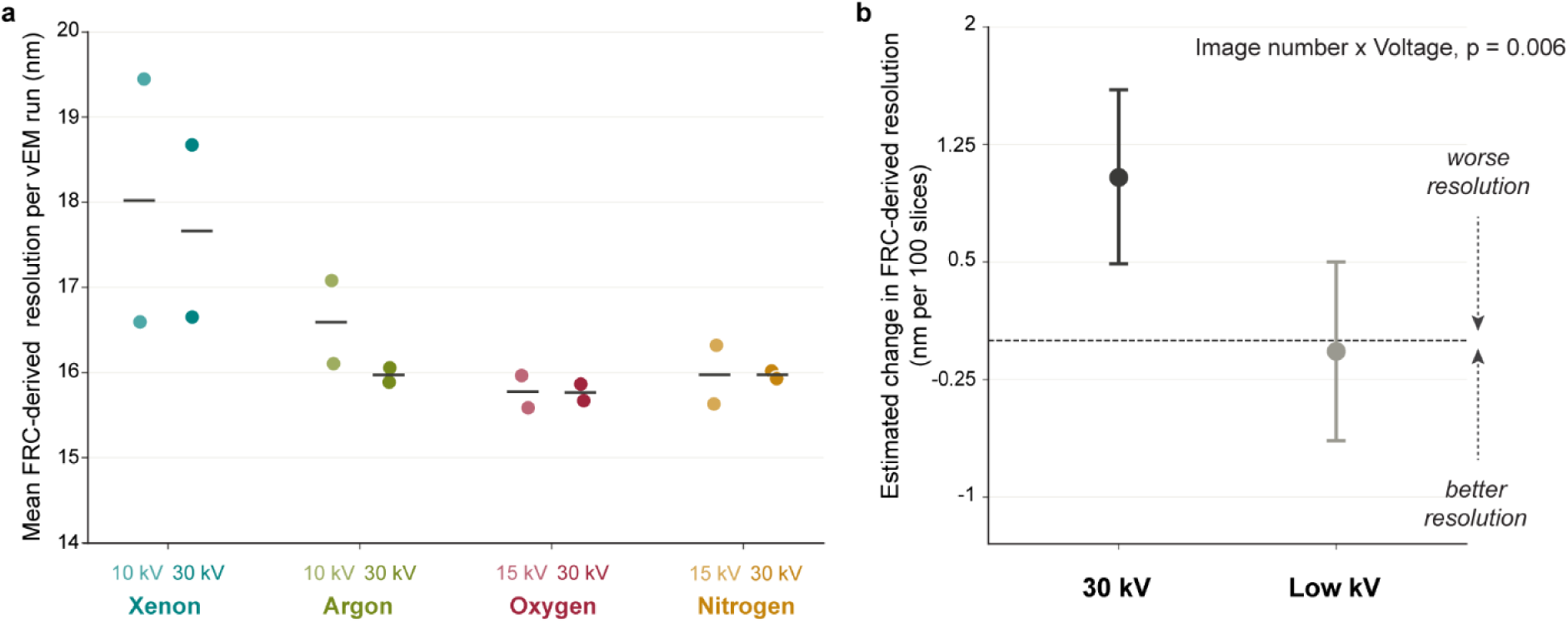
Mean FRC-derived SEM resolution during automated volume acquisition and across milling conditions. (A) Mean patch-based FRC-derived resolution of block-face SEM images acquired following milling with Xe, Ar, O and N at 30 kV and the corresponding species-specific low-kV conditions (10 kV for Xe and Ar; 15 kV for O and N). FRC-derived resolution was calculated from non-overlapping image patches intersecting the cellular region and calibrated against conventional two-image FRC measurements as described in Methods. Points represent mean value for each biological replicate and black horizontal lines represent combined mean value. (B) Temporal stability of FRC-derived resolution during automated cryo-pFIB-SEM volume acquisition in respect to voltage. Points show the model-estimated change in FRC-derived resolution per consecutive block-face images across high- and low-kV operating regimes, and error bars indicate 95% confidence intervals. Positive values indicate deterioration in spatial resolution. At 30 kV, FRC-derived resolution deteriorated by an estimated 1.04 nm per 100 images (95% CI, 0.49-1.60), whereas no systematic temporal change was detected across the species-specific low-kV conditions (−0.07 nm per 100 images; 95% CI, −0.64-0.50). Temporal behavior differed significantly between voltage regimes (image number × voltage interaction, p = 0.006; linear mixed-effects model). Individual acquisition trajectories are shown in Supplementary Fig 9.

**Supplementary Table 1.**
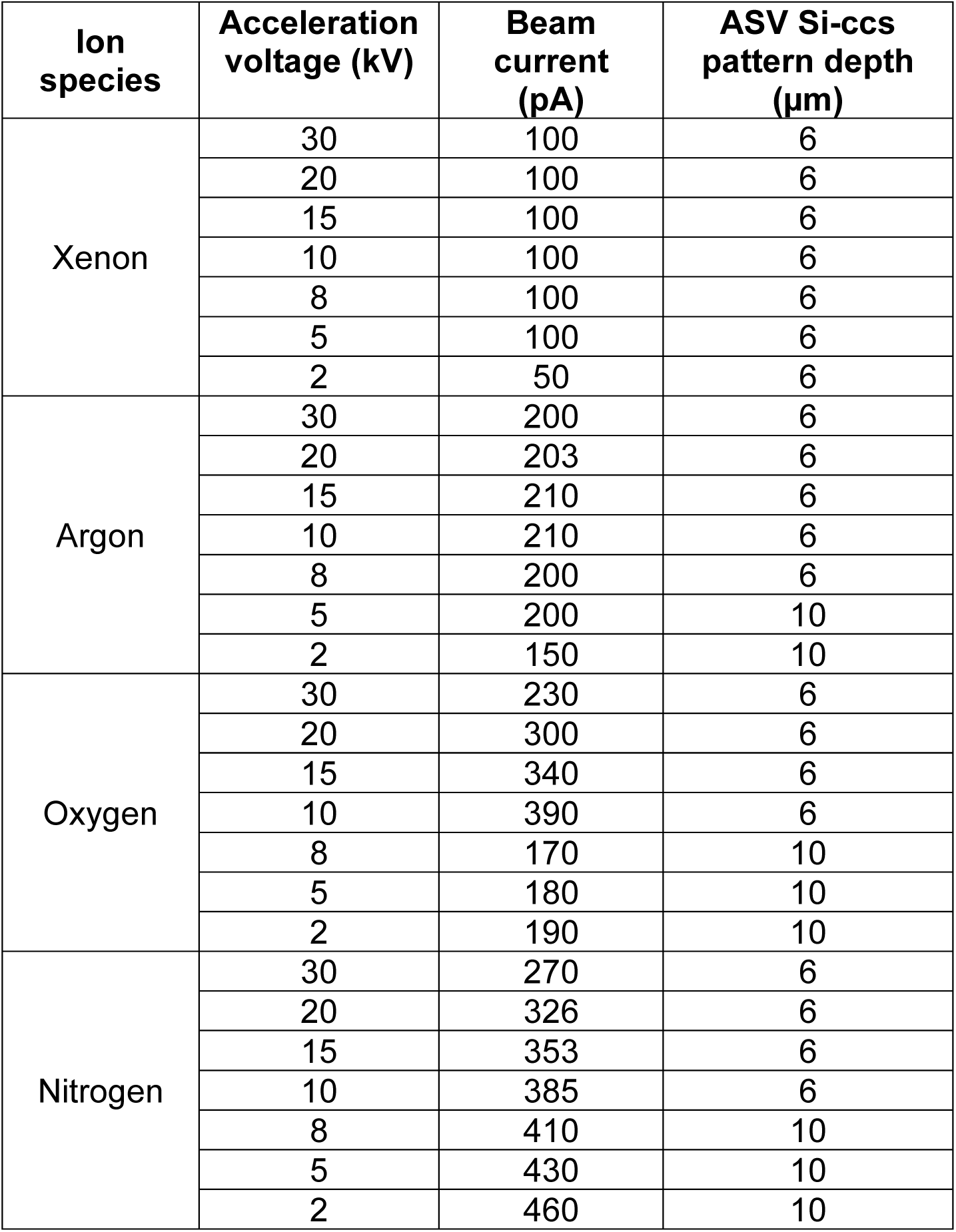
Operational pFIB ion beam currents on Hydra Bio UX when milling with specific voltages using specific ion species.

| **Ion species** | **Acceleration voltage (kV)** | **Beam current (pA)** | **ASV Si-ccs pattern depth (μm)** |
| --- | --- | --- | --- |
| Xenon | 30 | 100 | 6 |
|  | 20 | 100 | 6 |
|  | 15 | 100 | 6 |
|  | 10 | 100 | 6 |
|  | 8 | 100 | 6 |
|  | 5 | 100 | 6 |
|  | 2 | 50 | 6 |
| Argon | 30 | 200 | 6 |
|  | 20 | 203 | 6 |
|  | 15 | 210 | 6 |
|  | 10 | 210 | 6 |
|  | 8 | 200 | 6 |
|  | 5 | 200 | 10 |
|  | 2 | 150 | 10 |
| Oxygen | 30 | 230 | 6 |
|  | 20 | 300 | 6 |
|  | 15 | 340 | 6 |
|  | 10 | 390 | 6 |
|  | 8 | 170 | 10 |
|  | 5 | 180 | 10 |
|  | 2 | 190 | 10 |
| Nitrogen | 30 | 270 | 6 |
|  | 20 | 326 | 6 |
|  | 15 | 353 | 6 |
|  | 10 | 385 | 6 |
|  | 8 | 410 | 10 |
|  | 5 | 430 | 10 |
|  | 2 | 460 | 10 |

**Supplementary Table 2.** Imaging conditions used in the block-face damage assessment experiment.

| **Imaging** | **Detector,**<br>**mode** | **HFW**<br>**(pixel width)** | **Dwell** | **Line**<br>**integration** | **Contrast /**<br>**Brightness** |
| --- | --- | --- | --- | --- | --- |
| On-axis SE | TLD, DHV | 15.36 $\mu\text{m}$<br>(5.0 nm) | 50 ns | 150 | 77 / 23 |
| Off-axis SE | TLD, DHV | 7.68 $\mu\text{m}$<br>(2.5 nm) | 50 ns | 200 | 77 / 23 |
| On-axis BSE | TLD, BSE | 15.36 $\mu\text{m}$<br>(5.0 nm) | 4 $\mu\text{s}$ | 9 | 82 / 45 |

**Supplementary Table 3.**
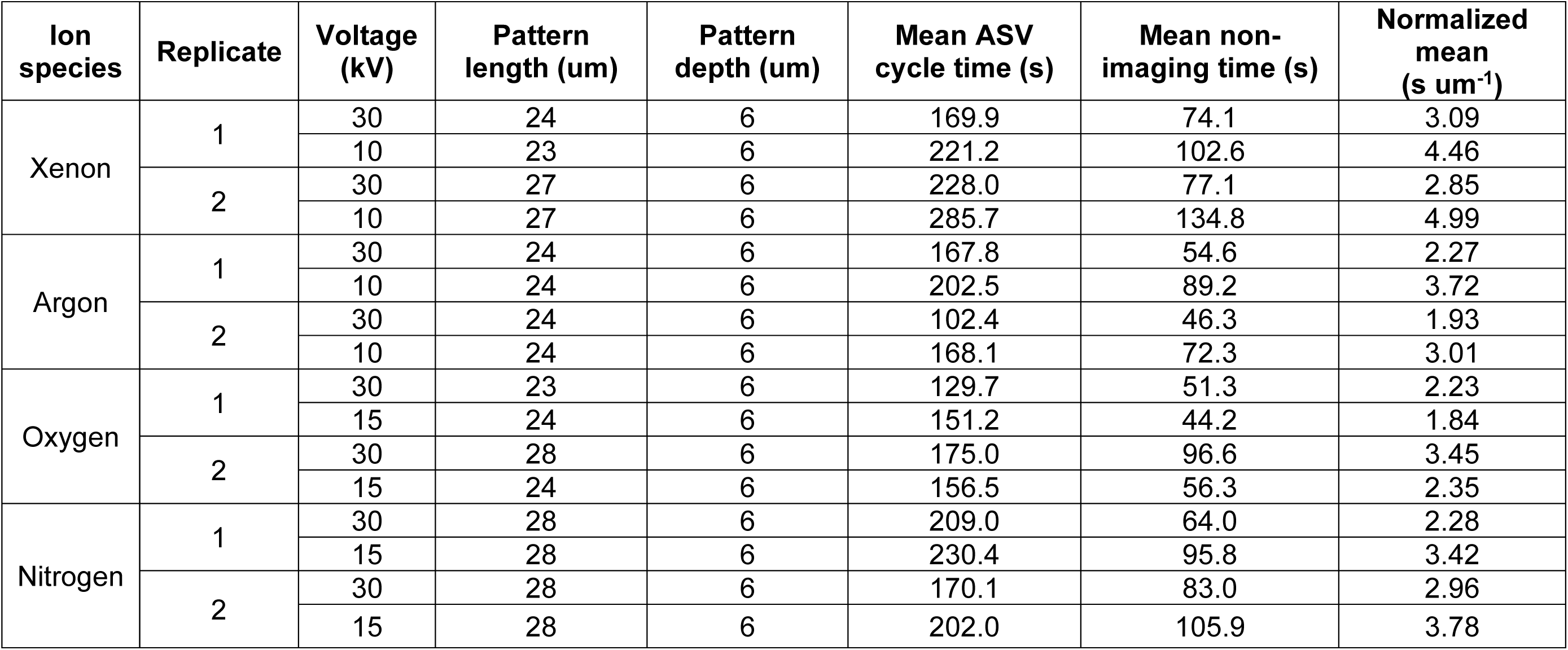
Acquisition time during automated cryo-pFIB-SEM volume imaging at high and reduced milling voltages. Mean non-imaging time was calculated after subtraction of the corresponding SEM frame acquisition time from each remaining slice-to-slice interval. To account for differences in the lateral extent of the milled regions, mean non-imaging time was normalized to the corresponding milling/pattern length and expressed as s µm⁻¹. High- and reduced-voltage conditions represent paired measurements from biological replicates.

**Supplementary Table 4.** The effect of reduced FIB accelerating voltage on cryo-pFIB-SEM volume-imaging acquisition time. Values show the percentage change in normalized mean non-imaging time at reduced accelerating voltage relative to 30 kV for each biological replicate. Positive values indicate an increase and negative values a decrease in normalized non-imaging time. Comparisons were performed within biological replicates. Reduced-voltage conditions were 10 kV for xenon and argon and 15 kV for oxygen and nitrogen.

| Ion species | Voltage | Replicate 1 | Replicate 2 | Milling time |
| --- | --- | --- | --- | --- |
| Xenon | 30 → 10 kV | +44.5% | +75.0% | Increase |
| Argon | 30 → 10 kV | +63.5% | +56.1% | Increase |
| Oxygen | 30 → 15 kV | -17.5% | -32.0% | Decrease |
| Nitrogen | 30 → 15 kV | +49.8% | +27.6% | Increase |

**Supplementary Video 1.** 3D visualization of a cryo-pFIB-SEM volume acquired while using xenon milling at 30 kV and the corresponding machine learning-based segmentation (biological replicate 1). The video shows the raw volume followed by segmentation of the nuclear membrane (blue), late endosomes (magenta), mitochondria (green), lipid bodies (orange) and charged regions (white).

**Supplementary Video 2.** 3D visualization of a cryo-pFIB-SEM volume acquired while using xenon milling at 30 kV and the corresponding machine learning-based segmentation (biological replicate 2). The video shows the raw volume followed by segmentation of the nuclear membrane (blue), late endosomes (magenta), mitochondria (green), Golgi apparatus (light blue), lipid bodies (orange) and charged regions (white).

**Supplementary Video 3.** 3D visualization of a cryo-pFIB-SEM volume acquired while using xenon milling at 10 kV and the corresponding machine learning-based segmentation (biological replicate 1). The video shows the raw volume followed by segmentation of the nuclear membrane (blue), late endosomes (magenta), mitochondria (green) and charged regions (white).

**Supplementary Video 4.** 3D visualization of a cryo-pFIB-SEM volume acquired while using xenon milling at 10 kV and the corresponding machine learning-based segmentation (biological replicate 2). The video shows the raw volume followed by segmentation of the late endosomes (magenta), mitochondria (green), Golgi apparatus (light blue) and charged regions (white).

**Supplementary Video 5.** 3D visualization of a cryo-pFIB-SEM volume acquired while using argon milling at 30 kV and the corresponding machine learning-based segmentation (biological replicate 1). The video shows the raw volume followed by segmentation of the nuclear membrane (blue), late endosomes (magenta), mitochondria (green), Golgi apparatus (light blue), lipid bodies (orange) and charged regions (white).

**Supplementary Video 6.** 3D visualization of a cryo-pFIB-SEM volume acquired while using argon milling at 30 kV and the corresponding machine learning-based segmentation (biological replicate 2). The video shows the raw volume followed by segmentation of the nuclear membrane (blue), late endosomes (magenta), mitochondria (green), Golgi apparatus (light blue), lipid bodies (orange) and charged regions (white).

**Supplementary Video 7.** 3D visualization of a cryo-pFIB-SEM volume acquired while using argon milling at 10 kV and the corresponding machine learning-based segmentation (biological replicate 1). The video shows the raw volume followed by segmentation of the nuclear membrane (blue), late endosomes (magenta), mitochondria (green), lipid bodies (orange) and charged regions (white).

**Supplementary Video 8.** 3D visualization of a cryo-pFIB-SEM volume acquired while using argon milling at 10 kV and the corresponding machine learning-based segmentation (biological replicate 2). The video shows the raw volume followed by segmentation of the nuclear membrane (blue), late endosomes (magenta), mitochondria (green), Golgi apparatus (light blue), lipid bodies (orange) and charged regions (white).

**Supplementary Video 9.** 3D visualization of a cryo-pFIB-SEM volume acquired while using oxygen milling at 30 kV and the corresponding machine learning-based segmentation (biological replicate 1). The video shows the raw volume followed by segmentation of the nuclear membrane (blue), late endosomes (magenta), mitochondria (green), Golgi apparatus (light blue), lipid bodies (orange) and charged regions (white).

**Supplementary Video 10.** 3D visualization of a cryo-pFIB-SEM volume acquired while using oxygen milling at 30 kV and the corresponding machine learning-based segmentation (biological replicate 2). The video shows the raw volume followed by segmentation of the nuclear membrane (blue), late endosomes (magenta), mitochondria (green), Golgi apparatus (light blue), lipid bodies (orange) and charged regions (white).

**Supplementary Video 11.** 3D visualization of a cryo-pFIB-SEM volume acquired while using oxygen milling at 15 kV and the corresponding machine learning-based segmentation (biological replicate 1). The video shows the raw volume followed by segmentation of the nuclear membrane (blue), late endosomes (magenta), mitochondria (green), Golgi apparatus (light blue), lipid bodies (orange) and charged regions (white).

**Supplementary Video 12.** 3D visualization of a cryo-pFIB-SEM volume acquired while using oxygen milling at 15 kV and the corresponding machine learning-based segmentation (biological replicate 2). The video shows the raw volume followed by segmentation of the nuclear membrane (blue), late endosomes (magenta), mitochondria (green) and charged regions (white).

**Supplementary Video 13.** 3D visualization of a cryo-pFIB-SEM volume acquired while using nitrogen milling at 30 kV and the corresponding machine learning-based segmentation (biological replicate 1). The video shows the raw volume followed by segmentation of the nuclear membrane (blue), late endosomes (magenta), mitochondria (green), Golgi apparatus (light blue), lipid bodies (orange) and charged regions (white).

**Supplementary Video 14.** 3D visualization of a cryo-pFIB-SEM volume acquired while using nitrogen milling at 30 kV and the corresponding machine learning-based segmentation (biological replicate 2). The video shows the raw volume followed by segmentation of the nuclear membrane (blue), late endosomes (magenta), mitochondria (green), lipid bodies (orange) and charged regions (white).

**Supplementary Video 15.** 3D visualization of a cryo-pFIB-SEM volume acquired while using nitrogen milling at 15 kV and the corresponding machine learning-based segmentation (biological replicate 1). The video shows the raw volume followed by segmentation of the nuclear membrane (blue), late endosomes (magenta), mitochondria (green), Golgi apparatus (light blue), lipid bodies (orange) and charged regions (white).

**Supplementary Video 16.** 3D visualization of a cryo-pFIB-SEM volume acquired while using nitrogen milling at 15 kV and the corresponding machine learning-based segmentation (biological replicate 2). The video shows the raw volume followed by segmentation of the nuclear membrane (blue), late endosomes (magenta), mitochondria (green), Golgi apparatus (light blue) and charged regions (white).

